# Modeling Friedreich’s ataxia with Bergmann glia-enriched human cerebellar organoids

**DOI:** 10.1101/2025.05.16.654315

**Authors:** Seungmi Ryu, Jason Inman, Hyenjong Hong, Vukasin M Jovanovic, Yeliz Gedik, Yogita Jethmalani, Inae Hur, Ty Voss, Justin Lack, Jack Collins, Pinar Ormanoglu, Anton Simeonov, Carlos A. Tristan, Ilyas Singeç

**Affiliations:** National Center for Advancing Translational Sciences (NCATS), Stem Cell Translation Laboratory (SCTL), National Institutes of Health (NIH), Rockville, MD 20850, USA; National Institute of Allergy and Infectious Diseases (NIAID), Collaborative Bioinformatics Resource (NCBR), National Institutes of Health (NIH), Bethesda, MD 20892, USA

## Abstract

The human cerebellum is a specialized brain region that is involved in various neurological and psychiatric diseases but has been challenging to study *in vitro* due its complex neurodevelopment and cellular diversity. Despite the progress in generating neural tissues from human induced pluripotent stem cells (iPSCs), an organoid model that recapitulates the key features of cerebellar development has not been widely established. Here, we report the generation of a 60-day method for human cerebellar organoids (hCBOs) that is characterized by induction of rhombomere 1 (R1) cellular identity followed by derivation of neuronal and glial cell types of the cerebellum. In contrast to forebrain organoids with multiple neural rosettes and inside-out neuronal migration, hCBOs develop a SOX2+ cerebellar plate on the outermost surface of organoids with outside-in neuronal migration, which is a characteristic hallmark of cerebellar histogenesis. These hCBOs produced various other cell types including granule neurons, Purkinje cells, Golgi neurons, and deep cerebellar nuclei. By using a glial induction strategy, we generate Bergmann glial cells (BGCs) within the hCBOs that not only serve as scaffolds for granule cells migration but also enhance electrophysiological response of the hCBOs. Furthermore, by generating hCBOs from patients with Friedreich’s ataxia (FRDA), we revealed abnormal disease-specific phenotypes that could be reversed by histone deacetylase (HDAC) inhibitors and gene editing by CRISPR-Cas9. Taken together, our advanced hCBO model provides new opportunities to investigate the molecular and cellular mechanisms of cerebellar ontogenesis and utilize patient-derived iPSCs for translational research.

## Introduction

The mammalian cerebellum is a unique brain region that plays important roles not only in motor coordination, balance, and posture but also in cognitive function (i.e., learning and memory) and processing of language and emotions.^1^ The highly complex anatomy and physiology of the human cerebellum can be affected in a broad range of early- and late-onset diseases with neurodevelopmental, genetic, and neurodegenerative causes (e.g., FRDA, Joubert syndrome, Dandy-Walker malformation, drug intoxication).^2^ In addition, the cerebellum is affected in neuropsychiatric disorders including autism spectrum disorders, schizophrenia, and depression.^3^

During the early stages of brain development, the cerebellum originates from R1, a defined region at the midbrain-hindbrain boundary that is caudal to the isthmic organizer.^4^ The R1 region gives rise to the cerebellar plate with two distinct neural precursor populations forming the rhombic lip (RL) and the ventricular zone (VZ) of the 4^th^ ventricle, which together will contribute to the diverse cell types of the cerebellum.^5^ The RL gives rise to all excitatory cells including granule cells, unipolar brush cells, the glutamatergic deep cerebellar nuclei.^5^ The VZ produces the Purkinje cells, Golgi cells, basket cells, and the GABAergic deep cerebellar nuclei neurons.^5^

One of the characteristic features of cerebellar development is that the dorsally located RL cells migrate tangentially and generate the external germinal layer (EGL), a transient layer of progenitors at the pial surface that produces the vast majority of granule cells (GCs), which then migrate in an outside-in direction to find their final destination in the internal granular layer (IGL) in the more mature cerebellum.^6^ The emergence of BGCs, a specialized unipolar cell type with long processes, that are important in guiding migratory GCs from the EGL toward the IGL.^7^ Disruption of the unique migration processes in the developing cerebellum have been linked to neurodevelopmental disorders, cerebellar ataxias, and impaired motor function.^8, 9^

Human cerebellar development and the many diseases that affect the cerebellum, particularly rare diseases, remain largely understudied. Reliable *in vitro* systems that faithfully recapitulate the key aspects of human cerebellar development are urgently needed for basic and translational research as substantial genetic, anatomic, and physiological differences exist between humans and animal models.^1, 10, 11, 12, 13, 14, 15^ Here, we developed a new 60-day method for deriving hCBO from iPSCs exhibiting critical features of cerebellar development. To the best of our knowledge, hCBOs with clear formation of the cerebellar plate on the organoid surface, enriched Bergmann glia, followed by outside-in migration of granule cells has not previously been established. Furthermore, we generated hCBOs from patients with FRDA, the most common inherited ataxia caused by abnormal GAA triplet-repeats in the FXN gene that affects 1 in 50,000 people globally.^16, 17^ The use of patient-derived iPSC lines with short and long GAA repeats, which differentially impact frataxin expression levels and correlate with disease severity in the clinic, enabled the identification of disease-phenotypes that were reversed by chemical and genetic strategies.

## Results

### A cerebellar organoid model with outside-in migration

Considering the unique features of cerebellar development and guided by the principles of early patterning along the neuraxis (dorso-ventral and anterior-posterior), we developed a step-wise 60-day differentiation protocol that consists of four stages characterized by neural induction and R1 specification, formation of the cerebellar plate, emergence of Bergmann glia, and organoid maturation (Figure 1A). First, human pluripotent stem cell (hPSC) cultures were dissociated, and a defined number of cells were aggregated to form embryoid bodies (EBs) in growth factor-free chemically defined medium supplemented with the CEPT small molecule (Figure 1A).^10^ We previously demonstrated that the CEPT cocktail is essential for stress-free passaging of hPSCs and for promoting optimal aggregation of hPSCs into EBs.^18, 19^ The first stage of the protocol (day 0-14) was aimed at neural induction and patterning into R1, which gives rise to the entire cerebellum during fetal brain development. Of note, R1 is the most anterior segment of the hindbrain and the only hindbrain region that does not express HOX genes.^20^ Activating the WNT pathway has a caudalizing effect during early neural patterning and we performed dose-response experiments using the well-established WNT activator CHIR99021.^21^ This treatment induced the expression of WNT1 and FGF8, which are critical to the development of R1 (Figure 1B).^22, 23^ These experiments also revealed that the administration of 0.5 µM CHIR99021 was optimal to induce R1 identity, as demonstrated by the expression of transcription factors EN1 and EN2 (Figure 1C and S1A). Importantly, HOXA2, the most rostrally expressed HOX gene,^24^ was not induced at 0.5 µM CHIR99021 (Figure 1C), whereas higher concentrations of CHIR99021 (1 and 2 µM) induced enhanced caudalization as indicated by suppressed WNT1 and increased HOXA2 expression. Omitting CHIR99021 from the cultures showed lack of caudalization (Figure 1C).

**Figure 1.**
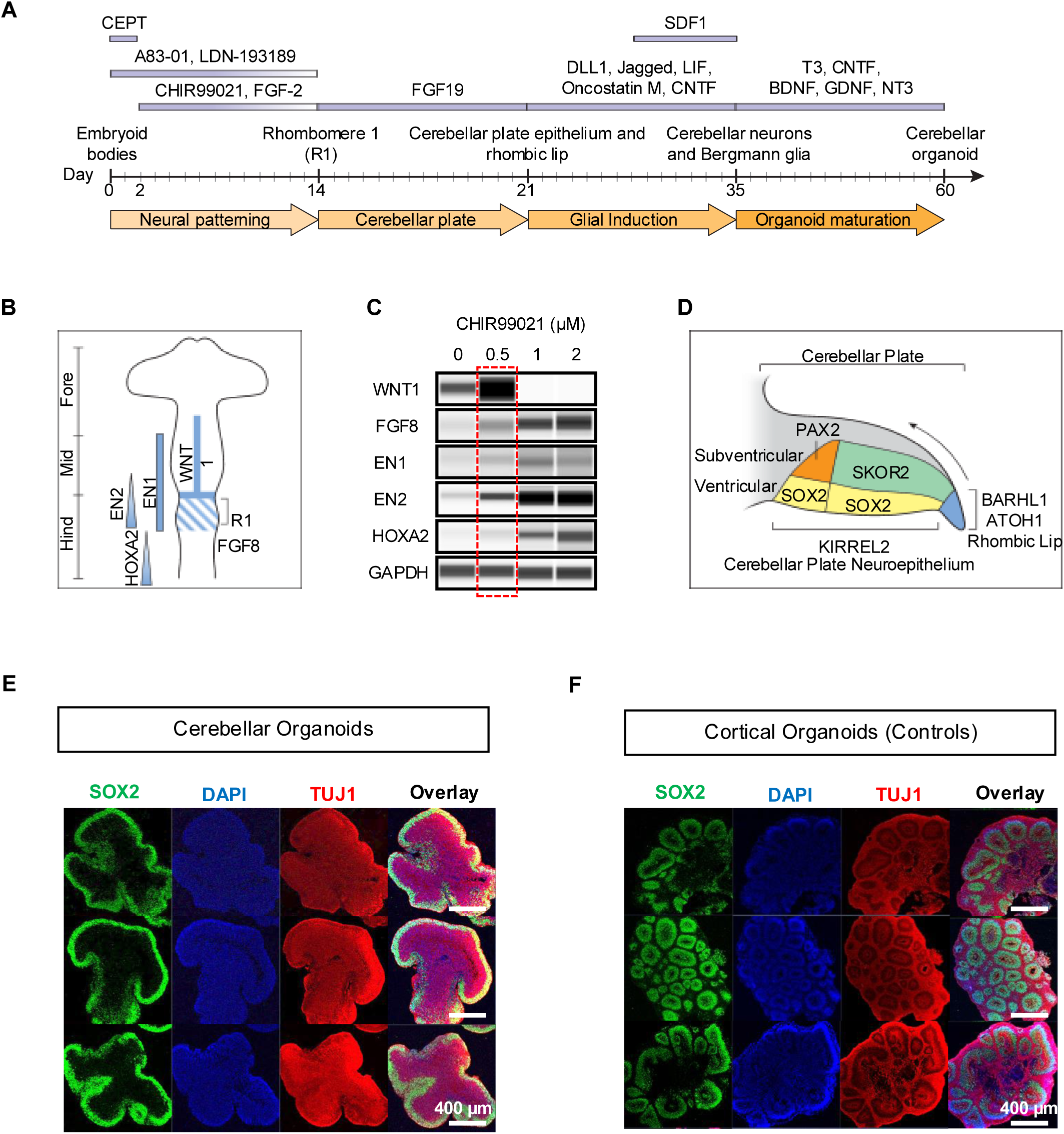
Establishment and characterization of hCBOs. (A) Overview of stepwise protocol for cerebellar organoid formation. (B) Gene expression pattern during regional specification of the embryonic brain. The cerebellum rises from the R1 region caudally adjacent to the mid-hindbrain boundary. (C) R1 patterning optimization via WNT activator CHIR99021 dose-response experiments in day 14 organoids. The red box indicates the optimal concentration for generating the R1 signature. (D) Schematic diagram showing the ventricular zone (VZ) and subventricular zone (SVZ) of cerebellar primordia, including the cerebellar plate neuroepithelium (CPNE) and the rhombic lip (RL). (E) Overview of three representative examples of iPSC-derived hCBOs (GM25256, day 21). Note the distinct formation of VZ (SOX2-expressing cells) on the organoid surface. Neuronal cells were immunostained using antibody against TUJ1. (F) Representative images of iPSC-derived cortical organoids (GM25256, day 35) with multiple SOX2+ expressing neural rosettes inside the organoids.

After inducing R1-like cellular identity, in the second stage of our protocol (day 14-21) cells were specified to generate the cerebellar plate by exposure to FGF19 (Figure 1A). The cerebellar plate contains the two primordial regions of the early developing cerebellum, the cerebellar plate neuroepithelium (CPNE) and the RL. The cells of these subregions express distinct markers including PAX2, SKOR2, KIRREL2 for subventricular zone (SVZ) cells originating from the VZ and BARHL1 and ATOH1 for the RL (Figure 1D).^10^ Importantly, immunocytochemical analysis of hCBOs at day 21 showed a histotypical cerebellar organization with SOX2+ precursors located exclusively on the organoid surface resembling the cerebellar primordium. In contrast to the cortical plate that generates the neocortex where neurons migrate from the inner layer to the outer layer, the cerebellar neurons follow an outside-in migratory pattern to find their final destination. Notably, SOX2+ cells were completely absent from the inner portion of hCBOs, which instead contained TUJ1+ neuronal cells (Figure 1E, S1A, S1B). To directly compare the difference between cortical and cerebellar organoids, in control experiments we generated cortical organoids following an established protocol.^25, 26^ In contrast to hCBOs, cortical organoids generated SOX2+ precursor cells in the inner region and TUJ1+ neurons on the outer region of neural rosetttes (Figure 1F). Hence, this direct comparison demonstrated that hCBOs with a distinct architecture resembling the early developing cerebellum can be derived using our protocol.

### Generation of Bergmann glia and the diversity of neuronal cell types of the cerebellum

Normal development and the proper formation of the cerebellar cortex depends on BGCs with long radial processes that serve as scaffolds for migratory granule cells.^6^ Because the spontaneous emergence of glial cells in organoids is a protracted process and can take 3-6 months,^25, 27^ in the third stage of our protocol (day 21-35) we took advantage of a combination of glial-inducing factors (Figure 1A) that we previously utilized as part of a different protocol to efficiently generate radial glial and astrocytes in a monolayer model.^28^ We also added the chemokine stromal-derived factor 1 (SDF1), which has important roles in regulating granule cell migration.^29^ This efficiently produced glial cells with long radial processes resembling Bergmann glia and showed expression of typical glial markers by day 35 including S100B, ALDH1L1, EAAT1, SOX9 and eventually abundant expression of GFAP at day 60 hCBOs (Figure 2A-2C).

**Figure 2.**
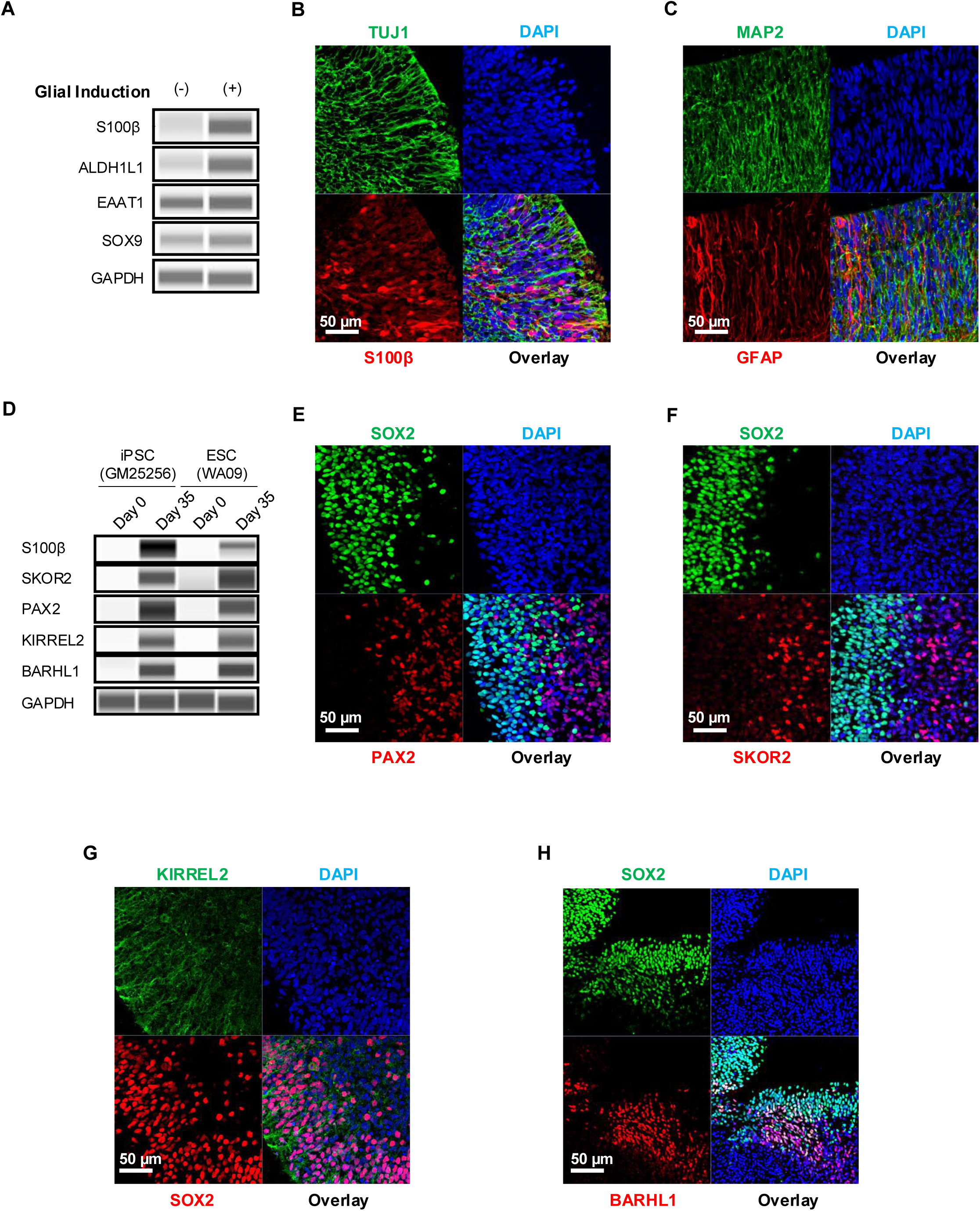
Generation and characterization of Bergmann glia and other cerebellar progenitors. (A) Western blot analysis showing the effects of glial induction on upregulation of glial markers. (B, C) Glial induction results in prominent expression of S100ß at day 35 and GFAP at day 60 hCBOs. Note the long radial processes of BGCs that are immunopositive for GFAP. (D) Western blot analysis showing expression of typical markers express by cells originating from the CPNE (S100Sß, SKOR2, PAX2, KIRREL2) or RL (BARHL1). (E, F) Immunostaining showing formation of CPNE. Note the distinct boundary between SOX2+ VZ and PAX2+, SKOR2+ SVZ. (G) Early purkinje cell marker KIRREL2 stains long processes extending into SVZ from SOX2-expressing neural precursor cells. (H) Immunostaining showing formation of RL. SOX2+ cells are intermingled and adjacent to cells expressing the RL marker BARHL1.

By day 35, in addition to SOX2+ precursors, distinct subpopulations of neuronal progenitors were detected via Western blots and immunohistochemical analyses (Figure 2D-2F, Figure S1A-C; see also schematic in Figure 1D). Purkinje cell progenitors expressing KIRREL2+ were also detected (Figure 2G). Cells expressing the CPNE marker (SKOR2, PAX2) and RL marker (PAX6, BARHL1, and ATOH1) were found adjacent to SOX2+ precursor cells (Figure 2H, Figure S1C). The overall data confirmed the presence of two primordial regions of the early developing cerebellum (CPNE and RL) by day 35 in hCBOs.

In the final step of our protocol (day 35-60), we matured hCBOs by adding a cocktail of neurotrophic factors and T3 thyroid hormone (Figure 1A). By day 60, Western blots and immunohistochemical analyses confirmed the expression of cell type-specific markers (Figure 3A-3H). Purkinje cells are GABAergic neurons and express the calcium-binding protein calbindin.^10,13^ We observed immature Purkinje neuron-like cells with various morphologies and immunoreactivity for calbindin and GABA (Figure 3B-3F). The neuronal protein NeuN is expressed by postmitotic cerebellar granule cells but not in Purkinje cells.^30^ Accordingly, we found a large number of NeuN+ granule cells that were devoid of calbindin expression (Figure 3F). Neurogranin is expressed by cerebellar Golgi cells and we found a subset of neurons expressing this marker (Figure 3G).^31^ Neurons of the deep cerebellar nuclei co-express TBR1 and SMI32, and cells expressing these markers were detected in hCBOs (Figure 3H).^10, 32^ Taken together, these data demonstrate that our 60-day protocol successfully generates hCBOs containing various types of cerebellar neurons and BGCs.

**Figure 3.**
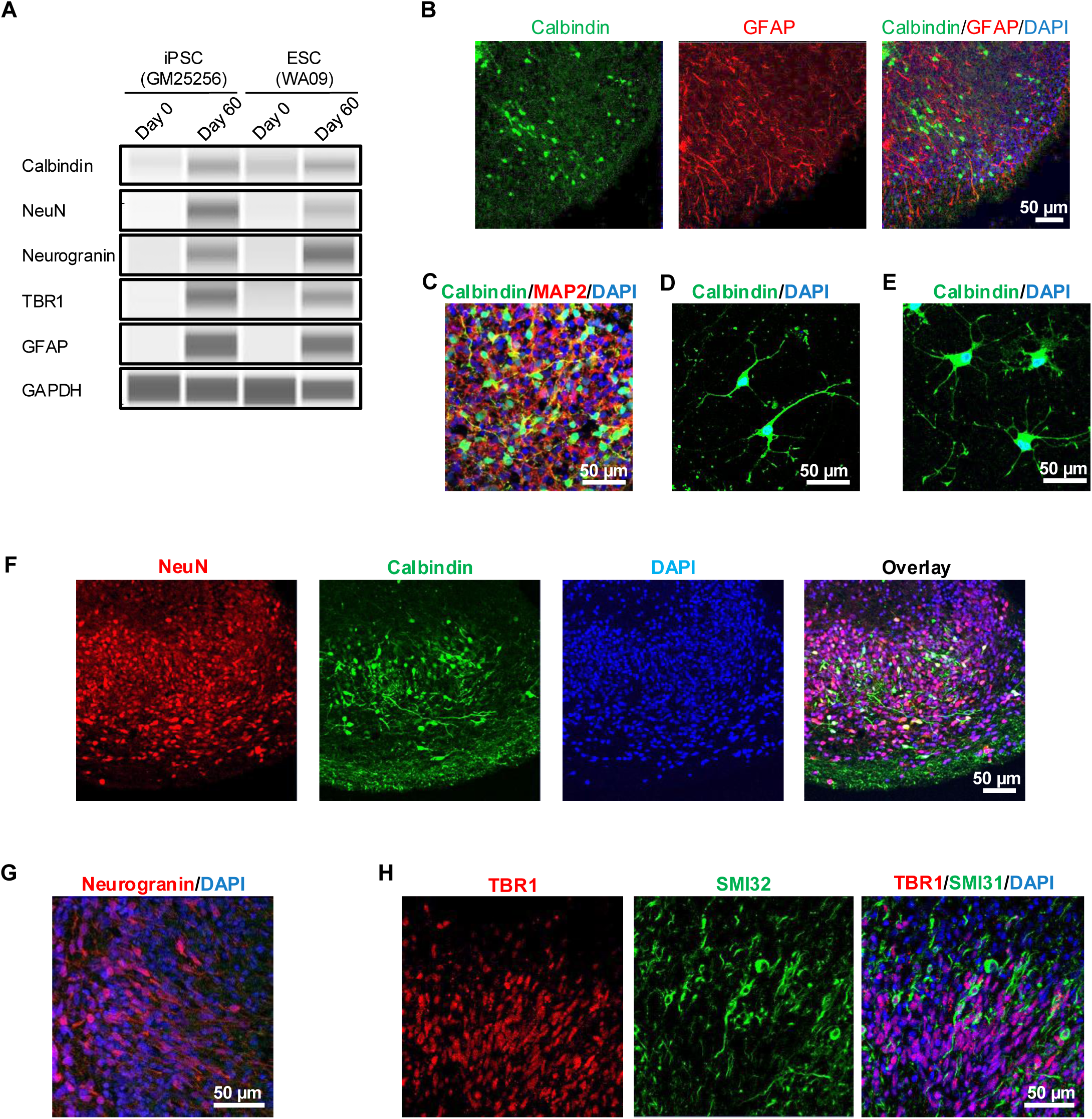
Immunophenotypic characterization of cerebellar neurons (day 60) (A) Western blot analysis demonstrating robust expressions of various cell type-specific cerebellar markers in hCBOs derived from an iPSC line (GM25256) and human ESC line (WA09). (B) Expression of calbindin by Purkinje cells and expression of GFAP by BGCs. (C) Densely packed neuronal cells expressing calbindin and neuronal marker MAP2. (D, E) Purkinje cells expressing calbindin observed in 2D tissue culture 30 days after dissociation of day-60 hCBOs. Note the complex neuronal morphologies of early Purkinje cells. (F) Visualization of large numbers of migratory GC (NeuN) and Purkinje cells (calbindin). (G) Expression of neurogranin by Golgi cells. (H) Immunostaining for TBR1 and SMI32 labeling neurons of deep cerebellar nuclei.

### Structural and functional consequences of glial induction in hCBOs

To characterize the role of Bergmann glia-like cells in the context of neuronal migration, we developed an image-based machine-learning approach to measure cell positioning of Purkinje neurons, granule cells, and BGC, respectively (Figure 4A-4B, Figure S2). When hCBOs were generated either with or without glial induction and subjected to quantitative morphological analysis, a marked difference was observed between the two groups. Organoids exposed to glial-inducing factors showed more prominent GFAP+ BGCs with long processes covering a larger area as compared to non-treated hCBOs (Figure 4A). Accordingly, NeuN+ granule cells and Calbindin+ Purkinje neurons were spread over a larger area suggesting more efficient outside-in migration guided by Bergmann glia scaffolds (Figure 4A-4B).

**Figure 4.**
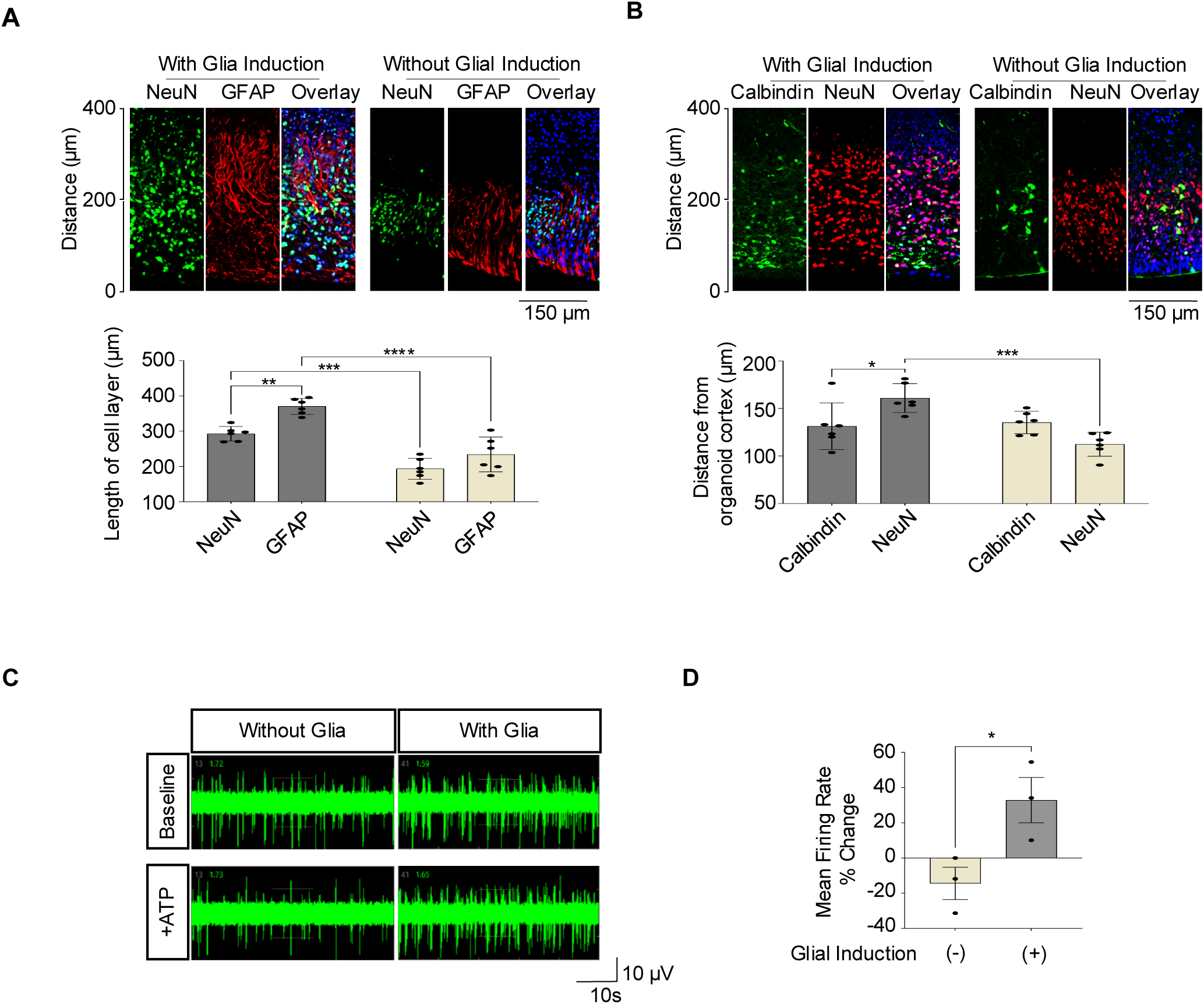
Glial induction and functional effects of BGCs. (A, B) Comparison of hCBOs with and without glial induction quantification of neuronal migration. Granule cells were visualized by using antibody against NeuN, Purkinje cells detected by calbindin expression, and BGCs by GFAP expression. Note the positive effect of glial induction and the presence of BGCs on layer formation and more widespread neuronal migration. n =3. Error bar, mean ± SEM. \**p*<0.05. \*\**p*<0.01. \*\*\**p*<0.001. \*\*\*\**p*<0.0001. (C, D) MEA analysis of hCBOs (day 80) stimulated with 100 uM ATP. n=3. Error bar, mean ± SEM. \**p*<0.05.

The presence of BGCs can modulate the electrophysiological activity of cerebellar neurons including Purkinje cells upon ATP stimulation.^33^ Microelectrode array (MEA) analysis of day 80 organoids demonstrated that while both organoids, either with or without glial induction, exhibited spontaneous electrical activity, significant changes in firing rate after the addition of ATP were only observed in hCBOs that underwent glial induction (Figure 4C-4D). Overall, these findings demonstrated that the presence of Bergmann glia-like cells in hCBOs supported neuronal migration during early layer formation and electrophysiological response to biological stimuli such as ATP.

### Time-course transcriptomic analysis of hCBOs from healthy donors and patients with FRDA

To characterize the differentiation of iPSCs into hCBOs, we performed RNA sequencing (RNA-seq) experiments at different timepoints (day 0, 14, 21, 35, 60) using three different cell lines, including two from healthy individuals (GM25256, NCRM5) and one patient donor with FRDA (GM23913). Time-course transcriptomic analysis of differentiating hCBOs and principal component analysis demonstrated clustering of samples according to the different developmental stages (Figure 5A), indicating reproducibility of the method and successful generation of hCBOs from healthy and patient-derived iPSC lines. Heatmap clustering analysis confirmed differentiation into R1 and cerebellar identity as measured by comparing differentially expressed (DE) genes at different time points (Figure 5B). As expected, pluripotency-associated genes POU5F1 (OCT4) and NANOG were expressed at day 0 and absent at day 14. Critical genes associated with the R1 region (e.g., WNT1, FGF8, EN1, and EN2) were expressed at day 14 and 21 (Figure 5B). At these timepoints, downregulation of midbrain marker OTX2 and upregulation of the hindbrain marker GBX2 suggested appropriate patterning into the caudal midbrain-hindbrain boundary where R1 arises (Figure 1B). By day 21 and 35, genes indicative of CPNE (PAX2, SKOR2, PTF1A) and RL (ATOH1, BARHL1) were strongly expressed in the hCBOs along with typical neuronal markers (NEUROD1, MAP2) and gliogenesis-associated genes (NFIA, SOX9). In more mature hCBOs (day 60), additional genes known to be upregulated in the developing cerebellum, such as CBLN1 (pre-cerebellin), LHX1 (transcription factor expressed by developing Purkinje cells), GAD1 (GABA synthesis), GRIN1 (NMDA receptor subunit), S100β and GFAP (BGC markers) were highly expressed (Figure 5B).

**Figure 5.**
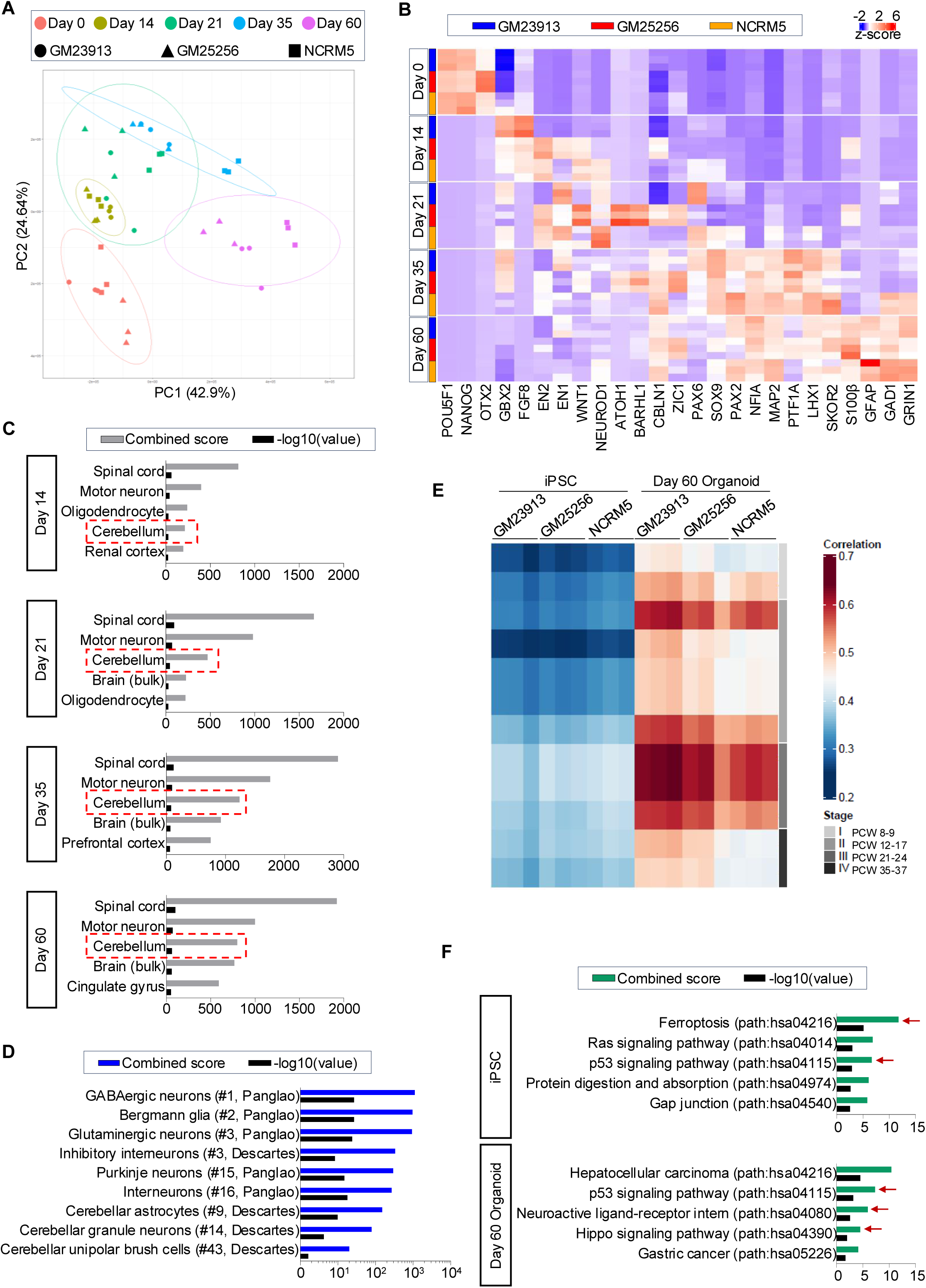
Transcriptomic analysis of hCBOs (day 60) (A) Principal component analysis plot of iPSC-derived hCBOs at each time point. Cerebellar organoids were derived from healthy donors (GM25256, NCRM5) and FRDA patient-derived (GM23913) iPSCs. (B) Heatmap analysis of key markers expressed at each time point during cerebellar organoid differentiation. n=3 independent hCBO batches at each time point for three independent cell lines. (C) Analysis comparing the transcriptome of cerebellar organoids at different time points to that of human tissue samples in the ARCHS4 database. Note that cerebellum was among the top hits across the different time points. (D) Comparison of transcriptomes of hCBOs to human cell atlases (Panglao and Descartes). (E) Heatmap of Spearman correlation analysis comparing day 60 hCBOs to fetal cerebellar tissues at different post-conception weeks (PCW, data from Allen BrainSpan). (F) KEGG pathway enrichment analysis reveals distinct pathway enrichment in iPSCs and cerebellar organoids from FRDA disease (GM23913) vs Healthy (GM25256, NCRM5) donors. Red arrows indicate metabolic pathways with relevance to FRDA pathophysiology. The top 500 differentially upregulated genes (adjusted P_val_ <0.05, Fold Change>2) were used in (C), (D) and (F).

To further confirm the transcriptomic signature of hCBOs, we performed unbiased gene enrichment analysis comparing the top 500 DE genes expressed by hCBOs to a publicly available tissue database containing 84,863 different human transcriptomes.^34^ This analysis underscored the transcriptomic signature of our organoids and “cerebellum” was a top-ranked category from day 14-60 (Figure 5C). Furthermore, we confirmed cerebellar cell type identities in hCBOs (day 60) by comparing the top DE genes with transcriptomic datasets from two large cellular databases, Panglao (>1 million cells analyzed from 74 tissues) and Descartes (>4 million cells analyzed from 15 tissues).^35, 36^ Again, this unbiased approach showed high correlations to various neuronal and glial cell types of the cerebellum (Figure 5D). The top categories included GABAergic/Glutamatergic neurons, Bergmann glia, and Purkinje cells, along with other cerebellar neuronal cell types, such as interneurons, granule neurons, and unipolar brush cells (Figure 5D). Next, to assess the maturity of hCBOs (day 60) relative to the developing cerebellum in the fetal brain, transcriptomic profiles were compared to publicly available datasets spanning from post-conception week (PCW) 8 to 37.^37^ This analysis revealed that hCBOs at day 60 were comparable to the fetal cerebellum at PCW 21-24 (Figure 5E).

Next, we asked if disease-associated changes could be detected in hCBOs from a FRDA patient (GM23913) when compared to the healthy controls (GM25256, NCRM5) (Figure 5F). Interestingly, gene set enrichment analysis showed upregulation in the p53 signaling pathway in FRDA samples in undifferentiated iPSCs as well as hCBOs (day 60). Upregulation of the p53 pathway was previously reported in the peripheral blood samples from 28 FRDA patients.^38^ Gene enrichment analysis also shed light on cancer pathways, ferroptosis, and Hippo signaling.

### Using hCBOs from FRDA patients to identify disease phenotypes

In FRDA, hyperexpansion of GAA repeats within the intron 1 region of the FXN locus induces repressive heterochromatin that is accompanied by hypoacetylation of histones H3 and H4.^39^ This hypoacetylation contributes to transcriptional silencing of the FXN gene leading to frataxin protein depletion within cells. Frataxin plays an important role in mitochondrial function as it controls iron-sulfur (Fe-S) cluster biogenesis within the mitochondria. Impairment of the iron-sulfur cluster machinery due to a depletion in frataxin leads to an increase in reactive oxygen species (ROS), followed by mitochondria dysfunction as shown through a decrease in mitochondrial enzyme aconitase 2 and eventually caspase-3 activation that causes cellular dysfunction and/or death (Figure 6A).^40^

**Figure 6.**
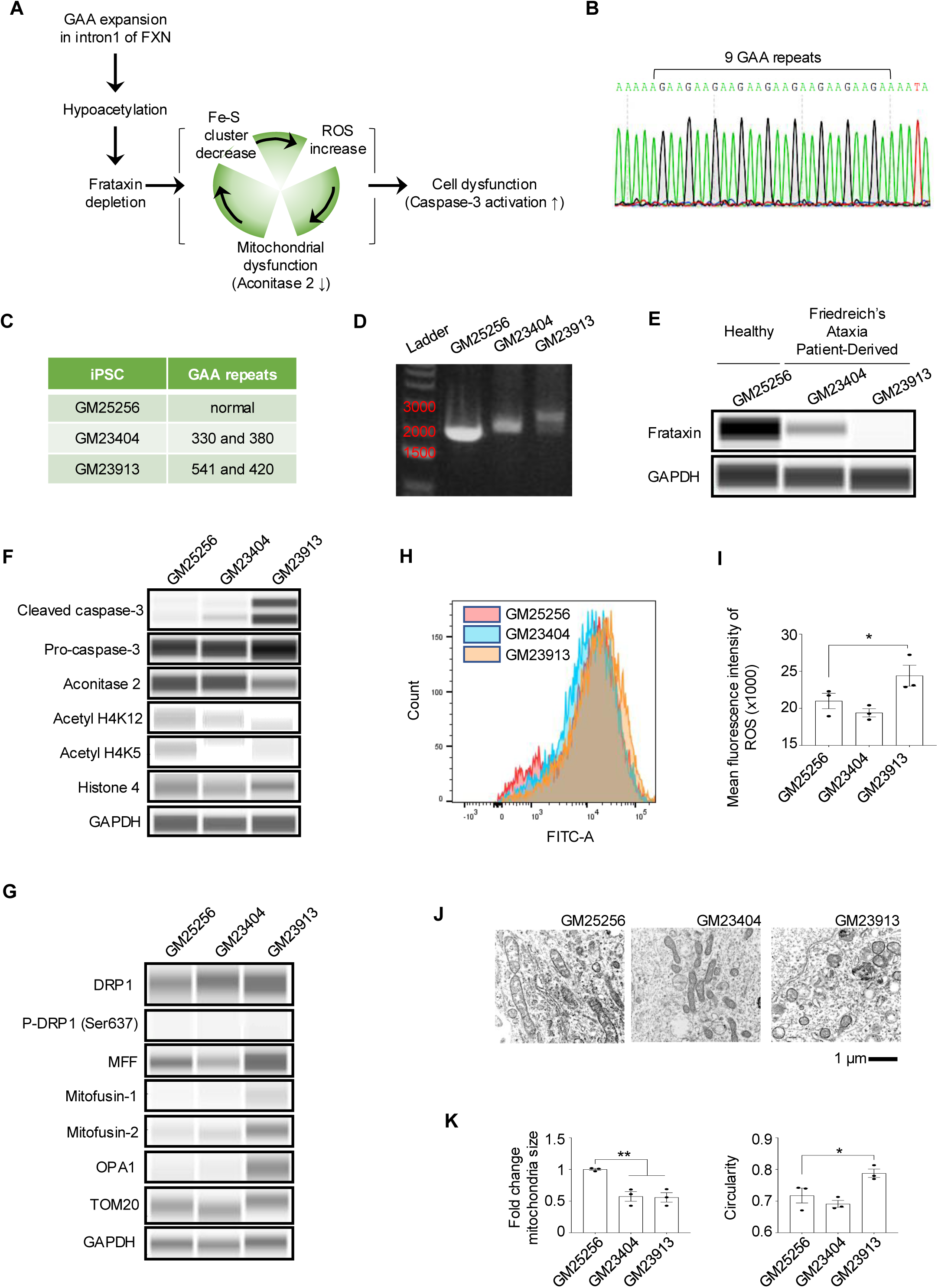
Generation of FRDA iPSC lines and identification of disease-specific phenotypes. (A) Schematic overview of FRDA pathology. Frataxin depletion results in impaired Fe-S (iron-sulfur) cluster synthesis and leads to increased ROS and mitochondrial dysfunction, ultimately resulting in cell death. (B) Sanger sequencing chromatogram showing GAA repeats within normal range in the Frataxin intron 1 region of GM25256 healthy iPSCs. (C) GAA repeat expansion lengths reported in Frataxin intron 1 locus of GM25256, GM23404, and GM23913 parental fibroblasts. (D) PCR genomic analysis of the Frataxin intron 1 confirming longer GAA expansion mutations in GM23404 and GM23913 iPSCs. (E) Western blot demonstrating decreased frataxin expression correlates with longer expansion mutations in FRDA iPSCs. (F) FRDA patient-derived hCBOs (day 60) exhibit increased levels of cleaved caspase-3, decreased mitochondrial enzyme aconitase 2, and hypo-acetylation of histone H4. Note that cerebellar organoids with longer GAA repeats show more distinct cellular pathology, which correlates with clinical disease severity. (G) Western blot analysis of mitochondrial proteins in hCBOs (day 60). Key proteins involved in mitochondrial fission (DRP1, P-DRP1, MFF), fusion (OPA1), and protein import (TOM20) were analyzed in healthy (GM25256) and FRDA (GM23404, GM23913) lines. (H, I) Quantification of intracellular ROS using fluorescence H2DCFDA demonstrates statistically significant elevation of ROS in day 60 FRDA hCBOs derived from the GM23913 line with the longer expansion mutation. n =3 biological replicates per group, \**p* < 0.05. (J, K) Electron microscopy images and quantification of mitochondrial size and circularity showing abnormal mitochondria in FRDA hCBOs (day 60). n = 3 images per group, \**p* < 0.05, \*\**p* < 0.01.

To demonstrate how hCBOs can be used for disease modeling, we performed a series of experiments that compared two FRDA patients-derived iPSC lines (GM23404 and GM23913) to a healthy control (GM25256). The iPSCs from both healthy and FRDA patients showed typical morphologies and expression of markers (OCT4, NANOG, and SOX2) associated with pluripotency (Figure S3A). Normal FXN alleles are known to have 6-34 GAA repeats,^16^ which was confirmed by Sanger sequencing in the healthy GM25256 iPSC line (9 GAA repeats, Figure 6B). Meanwhile, cell lines from FRDA patients had either 330/380 (GM23404) or 541/420 (GM23913) GAA repeats, respectively (Figure 6C). Furthermore, polymerase chain reaction (PCR) analysis with primers flanking the intron 1 region of FXN generated larger fragments in the FRDA iPSCs, confirming the presence of expansion mutations in lines GM23404 and GM23913 (Figure 6D). It is well-known that the length of the expansion mutation positively correlates with the extent of the FXN repression.^40^ Accordingly, in comparison to the healthy control (GM25256), patient-derived iPSCs with 330/380 GAA repeats (GM23404) had a reduction of FXN expression, while the iPSC line with 541/420 GAA repeats (GM23913) was completely devoid of FXN (Figure 6E).

Next, hCBOs were successfully generated from these three cell lines and expression of typical markers confirmed by immunohistochemistry and Western blot (Figure S3B-S3C). These organoids were composed of cells expressing typical CPNE markers (GBX2^+^, PAX2^+^, SKOR2^+^) and RL markers (PAX6^+^, BARHL1^+^) at day 35 (Figure S3B-S3C). When hCBOs were analyzed at day 60 by Western blot, various FRDA-associated disease phenotypes were detected including increased levels of activated/cleaved caspase 3, decreased level of aconitase 2 suggesting of mitochondrial dysfunction, and hypoacetylation of histone H4 (H4K5, H4K12) (Figure 6F). Of note, disease phenotypes were most severe in hCBOs from the patient with longer GAA expansion mutation (GM23913) as compared to healthy control and FRDA with a shorter expansion mutation (GM23404). The same trend and correlation with disease severity was observed when hCBOs were analyzed for expression of mitochondrial proteins implicated in mitochondrial function and plasticity (i.e., fission and fusion; Figure 6G) and ROS levels using flow cytometry (Figure 6H-6I). Lastly, ultrastructural analysis confirmed aberrant mitochondrial morphologies and differences in mitochondria size, which were more pronounced in the hCBOs generated from the iPSC line with longer GAA expansion mutations (Figure 6J-6K, S4).

### Correction of FRDA disease phenotypes

Having identified disease phenotypes characteristic of FRDA, we asked if it might be possible to correct these abnormalities. Previous studies tested whether frataxin expression could be induced in either FRDA patient-derived lymphocytes or iPSC-derived neurons in a 2D monolayer culture using HDAC inhibitors.^41, 42^ Using cerebellar organoids as the most relevant model for FRDA, we examined five commercially available HDAC inhibitors, testing three iPSC-derived hCBOs per drug (Figure 7A-7B). Among the tested compounds, trichostatin A (TSA) and valproic acid showed the strongest effects on inducing frataxin protein as measured by Western blot analysis. We also noted increased levels of H4 acetylation and marked reduction of activated caspase 3 (Figure 7A-7B). While HDAC inhibitors may have some beneficial effects, their long-term efficacy and safety for treating FRDA poses clinical challenges. For instance, the anti-epileptic drugs valproic acid, phenytoin, and others can damage the cerebellum and lead to loss of Purkinje cells and granule cells.^43, 44, 45^ Cell and gene therapies have emerged as therapeutic options as iPSCs and gene editing technologies have entered the clinical stage, while organoids can serve as predictive and physiologically-relevant preclinical models for genetic diseases.^46^ Establishment of gene-corrected isogenic iPSC lines using CRISPR-Cas9 has become an important strategy. Therefore, we employed a CRISPR-Cas9 gene editing system to excise the GAA expansion mutation in the FXN intron 1 region of FRDA patient-derived iPSC lines (Figure 7C) following a previous report.^40^ We confirmed via PCR gel-electrophoresis that the guide RNAs (gRNAs) resulted in effective excision of the GAA mutated region as shown through shorter bands detected at 152 bp (Figure 7D). To establish gene-corrected clonal iPSC lines, we used a single-cell printer and our previously established protocol including the CEPT small molecule cocktail to identify and expand new cell lines (Figure 7E).^18, 47^ Two clonal iPSC lines (clone 14 and 21) from the FRDA patient with longer GAA repeats (GM23913) were successfully established and characterized (Figures 7F, S5A-B). Long-read sequencing analysis (PacBio) verified on-target excision in the FXN intron 1 locus and the absence of off-target effects within the top four gRNA homology sites for gRNA1 and gRNA2 (Figure S5C-D, S6). Moreover, optical genomic mapping of the genomes confirmed normal karyotypes in the established clonal lines (Figure S7). Notably, increased frataxin expression was obtained in the isogenic gene-corrected FRDA iPSC lines (Figure 7G). By day 35, hCBOs showed normal development and differentiation into different cell types expressing typical markers as analyzed by Western blotting and immunohistochemistry (Figures 7H-I).

**Figure 7.**
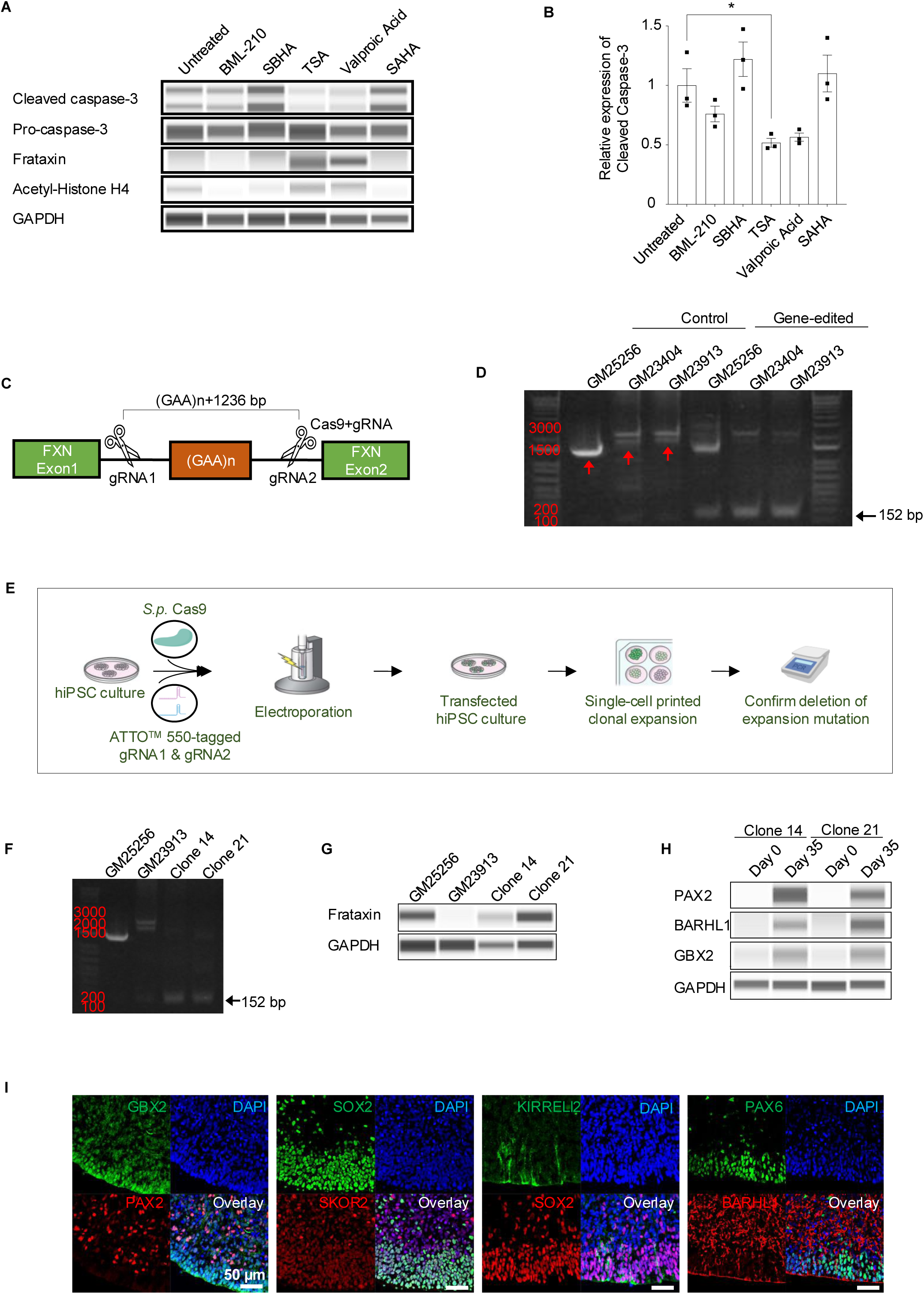
Drug testing in hCBOs from FRDA patients and gene-correction by using CRISPR-Cas9. (A) Effect of HDAC inhibitors on day 60 GM23913 hCBOs. SBHA: Suberohydroxamic acid. TSA: Trichostatin A. SAHA: Suberoylanilide hydroxamic acid. (B) Quantification of western blot analysis performed in (A). n = 3. Error bar, mean ± SEM, \**p*<0.05. (C) Gene editing strategy to correct the mutation in FRDA patient-derived iPSCs by using CRISPR-Cas9. gRNA: guide RNA (D) PCR analysis confirmed gene deletion in all the iPSC lines. The expected PCR product size post-gene editing is 152 bp. (E) Schematic overview depicting the strategy for generating clonal gene-corrected iPSC lines. (F) PCR analysis confirmed gene deletion in the established cell clones of corrected FRDA GM23913 iPSCs. (G) Western blot analysis of FXN expression in healthy (GM25256), disease (GM23913), and two gene-corrected isogenic lines from patient GM23913 (clone 14 and 21). (H, I) Western blot and immunohistochemical analysis (clone 14) of hCBOs (day 35) derived from gene-corrected isogenic GM23913 cell line.

## Discussion

Here, we established an advanced hCBO model that recapitulates key aspects of orchestrated cerebellar development including R1 specification, cerebellar plate formation, induction of Bergmann glia, outside-in migration, and derivation of the diversity of cerebellar cell types. Considering previous reports studying cerebellar organoids,^10, 13, 14, 15, 48^ to the best of our knowledge, this study is the first to demonstrate the generation of hCBOs enriched with functional BGCs that resulted in enhanced electrophysiology response and cellular migration within the hCBO. Our comparative transcriptomic analysis using RNA-seq confirmed stepwise cerebellar differentiation and also suggested that the molecular signature of hCBOs (days 60) was similar to the fetal cerebellum at PCW 21-24.

Our data demonstrated the generation of R1 cellular identity by using controlled WNT pathway activation. Similar to *in vivo* development, our approach promoted the expression of R1 markers (EN2, FGF8, WNT1), while hindbrain markers such as HOXA2 were not induced. In addition, utilizing our previously established combination of glial-inducing factors we efficiently generated BGCs within the hCBOs produced with this 60-day protocol.^28^ This approach underscores the fact that finding the optimal concentrations, and combination of small molecules and growth factors are critical for appropriate patterning and guiding cell differentiation trajectories.

Understanding the mechanisms by which BGCs contribute to cerebellum patterning and function has been of a long-standing interest in neuroscience.^49^ Abnormal BGC development is associated with various neurodevelopmental and genetic diseases,^8, 9^ yet availability of a complex human *in vitro* model has been limited until now. We predict that having access to cerebellar organoids, that can be generated within 60 days and produce BGCs, GCs and Purkinje cells, will be crucial for modeling and gaining a deeper understanding of cerebellar development and various diseases.

To demonstrate translational utility of hCBOs, we investigated iPSC lines from patients with FRDA. Consistent with the pathophysiology of FRDA and the known functions of the FXN gene and disease-severity correlating with the length of GAA repeats, we observed altered mitochondrial morphologies, oxidative stress, and elevated caspase 3 activation. The reversal of disease phenotypes by using HDAC inhibitors and generating gene-corrected isogenic cell lines by CRISPR-Cas9 was promising. Nevertheless, future studies are needed toward developing new therapeutic strategies that may include cell replacement and *in vivo* gene editing. We envisage that many other diseases affecting the cerebellum such as the recently described FGF14 GAA repeat expansion in late-onset cerebellar ataxia will benefit from investigating patient-derived hCBOs to advance the promise of precision medicine.^50^

### Limitations of the study

Our study significantly advances the use of hCBOs for fundamental research in developmental biology as well as disease modeling. However, we have not cultured hCBOs for extended periods of time, but it should be possible to extend culture period beyond 2-3 months as other groups have cultured organoids for 6-20 months.^13, 27^ It is likely that extended culture periods will result in increased morphological complexity and more mature cell types. This is particularly important for the generation and integration of Purkinje cells with highly complex dendritic arborizations, while their axons are the sole output from the cerebellar cortex. The identification and use of additional factors that promote neuronal maturation at these later stages could provide exciting opportunities to induce and study dendritic spines of Purkinje cells. Future experiments in more mature hCBOs could also uncover in more detail the spatial segregation, migratory pathways, and formation of the three-layered cerebellar cortex (i.e. molecular, Purkinje, and granular layer) in comparison to the deep cerebellar nuclei.

In hCBOs established from FRDA patients, many disease-specific phenotypes were previously reported and are consistent with our findings. Following our proof-of-principle experiments, testing a greater number of donor-derived iPSC lines will help to establish a powerful disease modeling platform to study various genotype-phenotype relationships and test new therapeutic modalities.

## Acknowledgments

Authors would like to thank Hannah Baskir, Juliana F. de Sousa, Glib Lirazan, Ewy A. Mathe, Dvir Blivis, the Automated Cell Technologies team at NCATS, Lara El Touny, and all the other colleagues at NCATS NIH for their support throughout this work. We would also like to thank NCI Frederick Core Facility, Vector Biolabs, VitroVivo Biotech, ACD Biotechne, and C M Cherry Consulting for their facility service support. We are grateful to Allan Hoofring and Ethan Tyler from the NIH Medical Arts Design Section for their artistic contribution. We also gratefully acknowledge funding from the Regenerative Medicine Program (RMP) of the NIH Common Fund and the intramural research program of the National Center for Advancing Translational Sciences (NCATS), NIH, grants TR000244 and TR000410. The funders had no role in study design, data collection, and analysis; decision to publish; or preparation of the manuscript.

## Author Contributions

S.R., and I.S. conceived the study and experiments. S.R., C.A.T., and I.S. worked on the experimental planning and data interpretation. S.R., J.I., H.H., V.M.J., Y.G., Y.J., P.O. performed the experiments. J.I., J.L., J.C. performed the bioinformatic analyses. I.H. T.V. performed the image analysis. S.R., J.I., H.H., V.M.J., Y.G., Y.J., I.H., T.V., V.J., J.L., J.C., P.O., A.S., C.A.T. and I.S. contributed to data analysis and discussion. S.R., C.A.T., and I.S. wrote, revised, and reviewed various drafts of the manuscript.

## Data Availability

Data available upon request.

## Figure Legends

**Figure S1.**
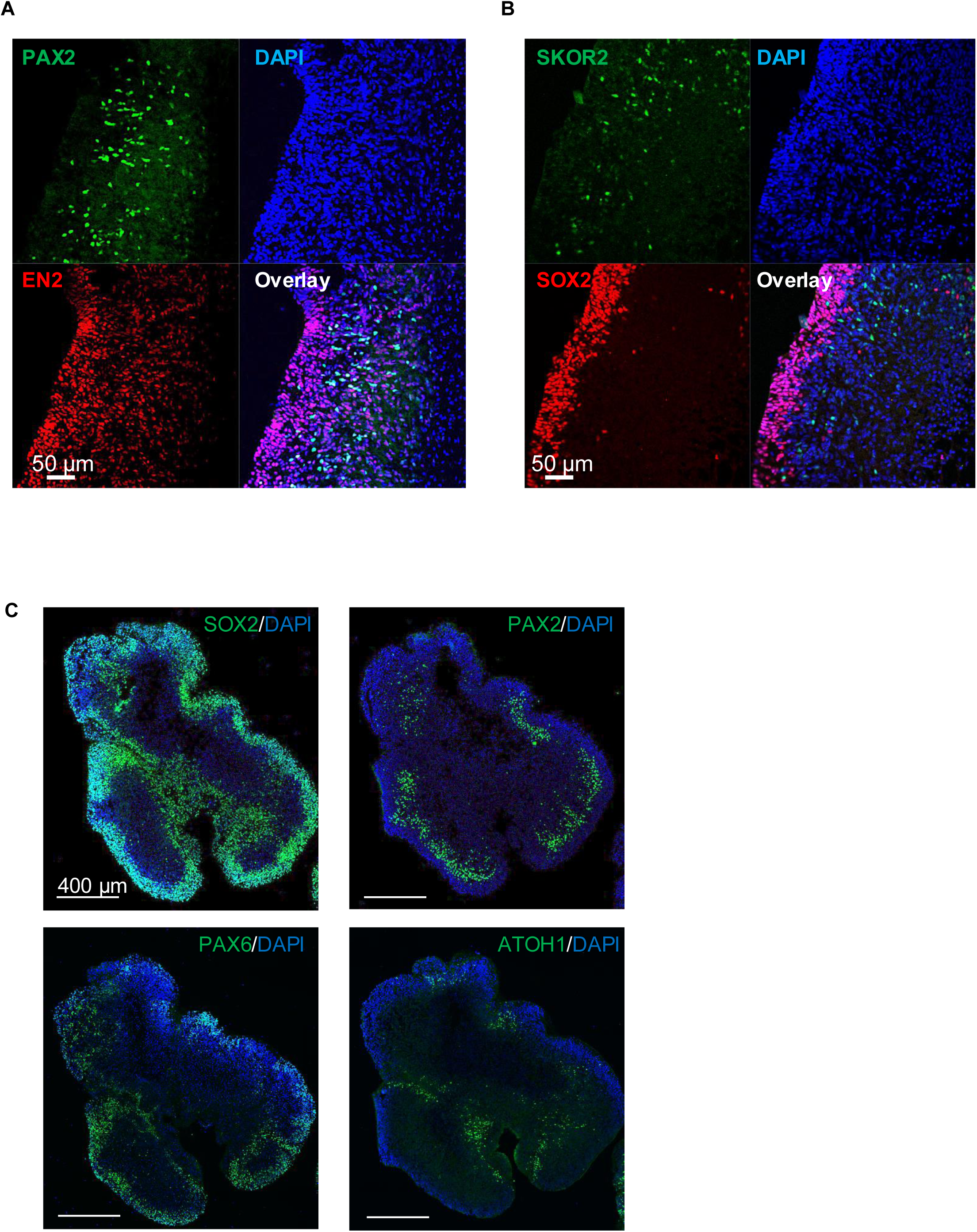
Immunohistochemical analysis of hCBOs. (A, B) Expression of typical markers of early CPNE development. Note the more widespread expression of R1 region marker EN2 along with the specification of VZ (SOX2) vs SVZ (PAX2, SKOR2). (C) Distribution of CPNE (PAX2) and RL (PAX6, ATOH1) across consecutive sections of the same organoid. All the data in (A)-(C) were obtained from day 35 hCBOs.

**Figure S2.**
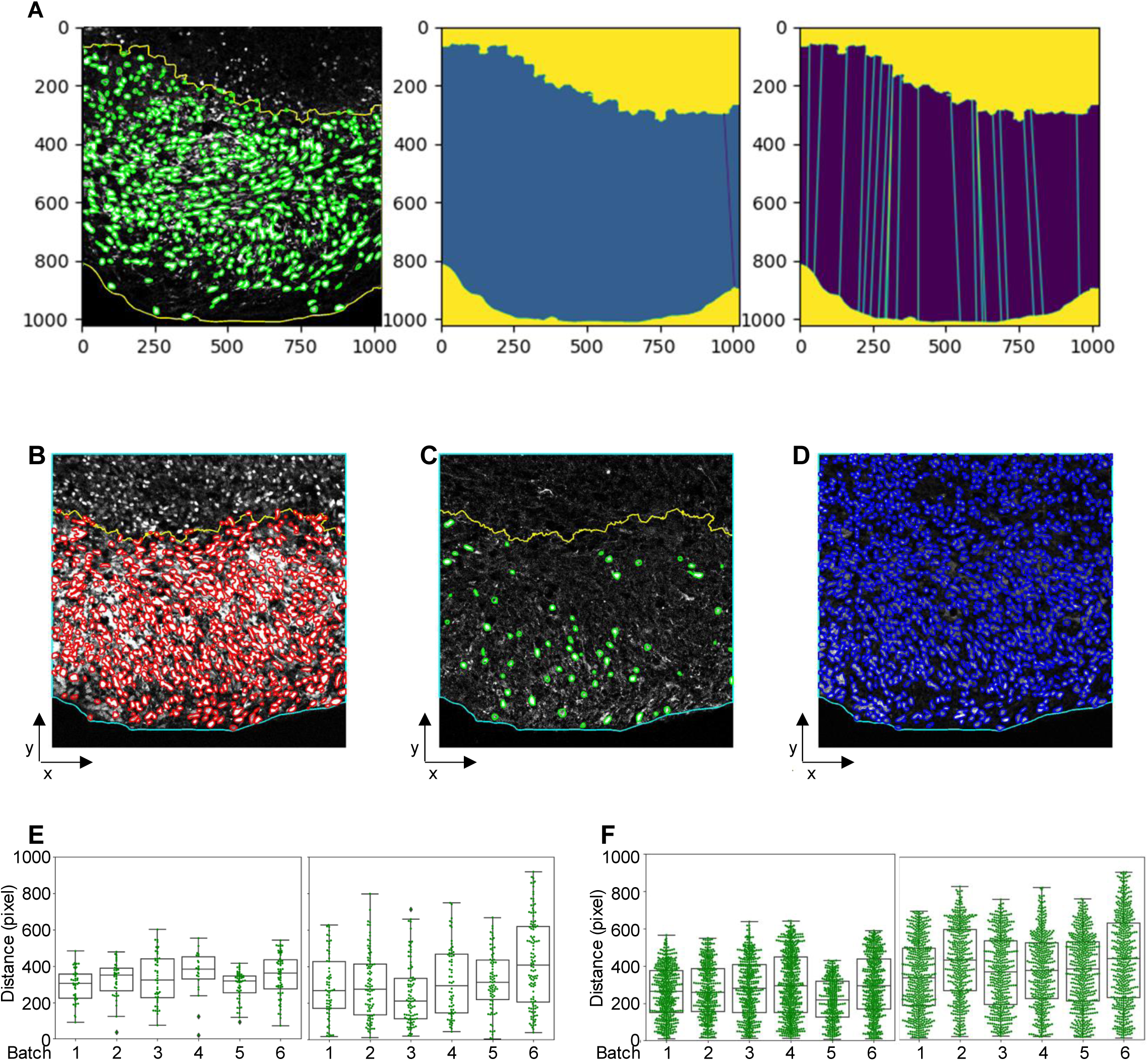
Machine learning-based image analysis to track neuronal migration. (A) Image-based analysis to quantify the length of the NeuN+ granule cell layer. The unit is shown in pixels. (B) Identification of calbindin+ Purkinje cells. (C) Identification of NeuN+ granule cells. (D) Identification of DAPI+ cell nuclei. (E) Box and Whisker plot showing the location of individual calbindin+ Purkinje cells relative to the cortex of the organoid. (F) Box and whisker plot showing the location of individual NeuN+ granule cells relative to the cortex of the organoid. The NeuN+ granule cells (masked green in (A) and red in (B)), calbindin+ Purkinje cells (masked green in (C)), and DAPI^+^ cell nuclei (masked blue in (D)) were identified using an optimized CellPose deep learning method. Upper and lower length regions containing NeuN+ cells were defined by the Super Pixel + Graph Cuts machine learning method and highlighted with yellow lines in (A) to quantify the length of the NeuN+ containing cell layer. Using the same method, the upper region of NeuN^+^ or calbindin^+^ cells (highlighted as yellow) and cortex of the organoid (highlighted as cyan lines) were defined in (B)-(D). 1 pixel equals 0.4151329 µm in (A), (E), and (F).

**Figure S3.**
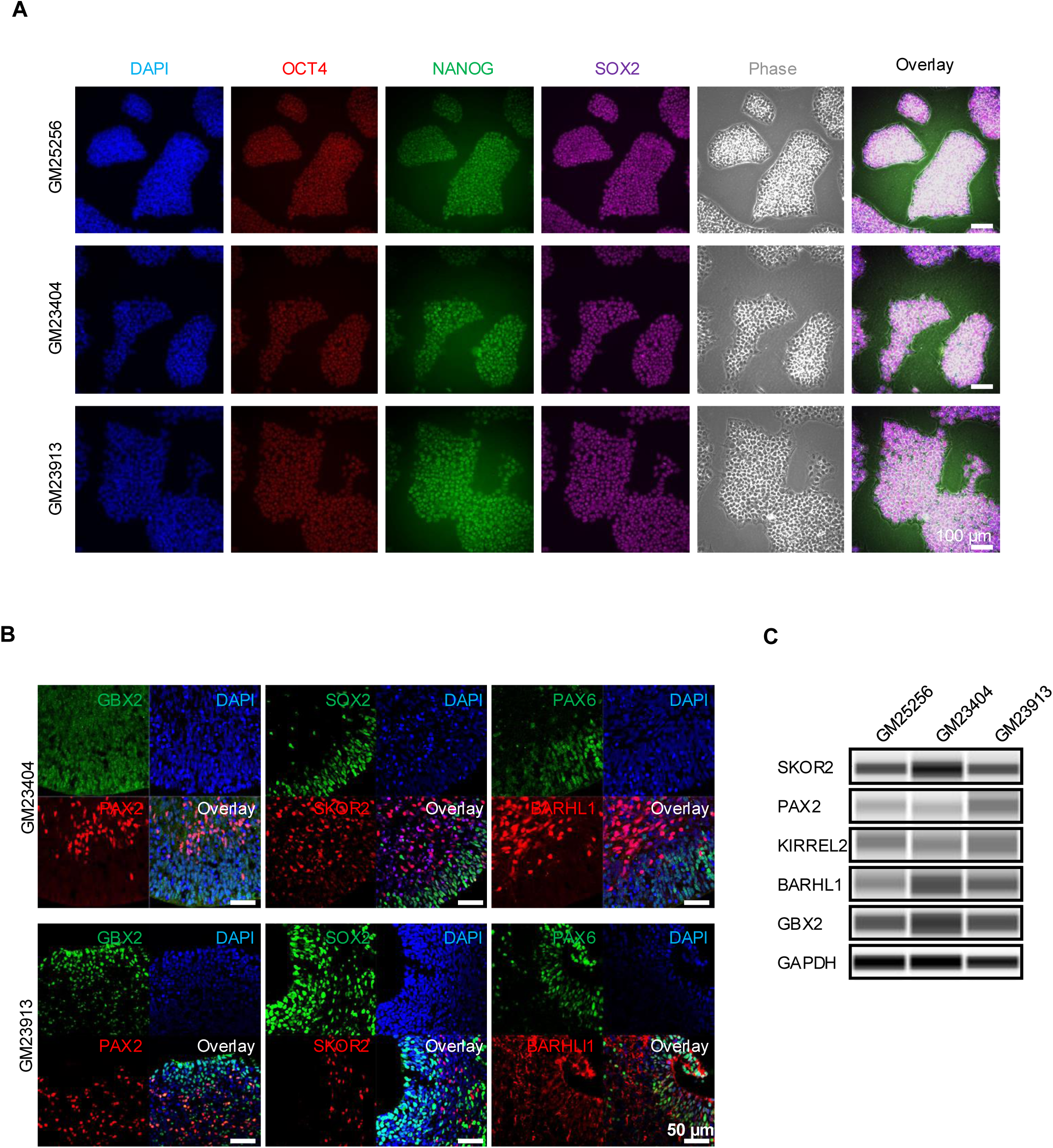
Characterization of iPSC lines and hCBOs from health and FRDA iPSC lines. (A) Expression of pluripotency-associated markers OCT4, NANOG, and SOX2 in iPSCs from a healthy individual (GM25256) and two FRDA patients. (B) Immunohistochemical analysis showing normal hCBO generation and expression of typical cerebellar markers at day 35. No apparent differences were observed between health and FRDA iPSC lines. (C) Western blot analysis showing primordial cerebellar markers (GBX2 for caudal region of mid-hindbrain boundary, BARHL1 for RL, and PAX2 of CPNE) expressed by corrected, isogenic GM23913 iPSC clones at day 35 compared to day 0.

**Figure S4.**
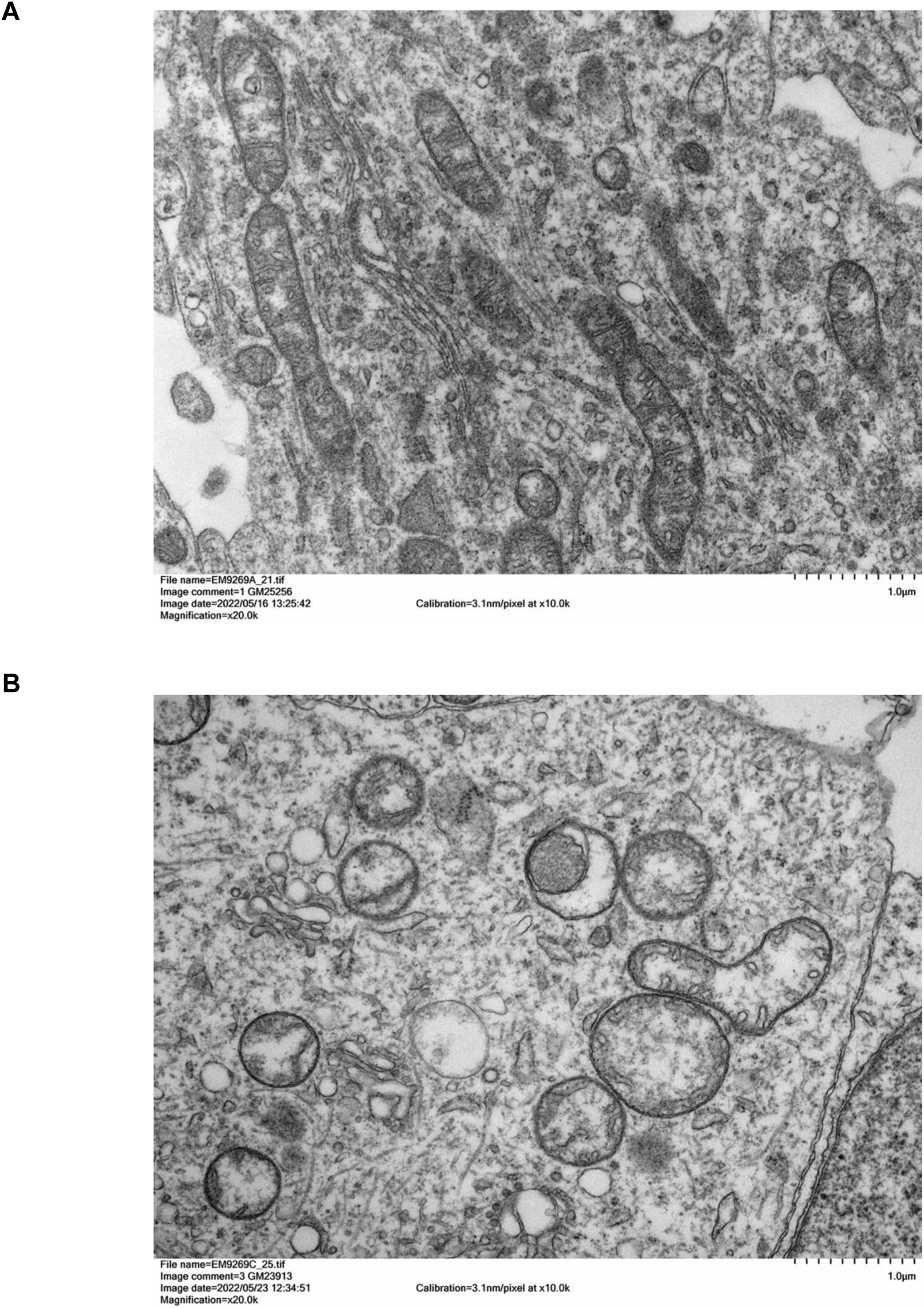
Ultrastructural analysis of mitochondria in normal and FRDA iPSC lines. (A, B) Higher magnification of electron microscopic images showing normal mitochondria (A) and mitochondria with pathological changes in FRDA (B). See also Figure 6J and 6K.

**Figure S5.**
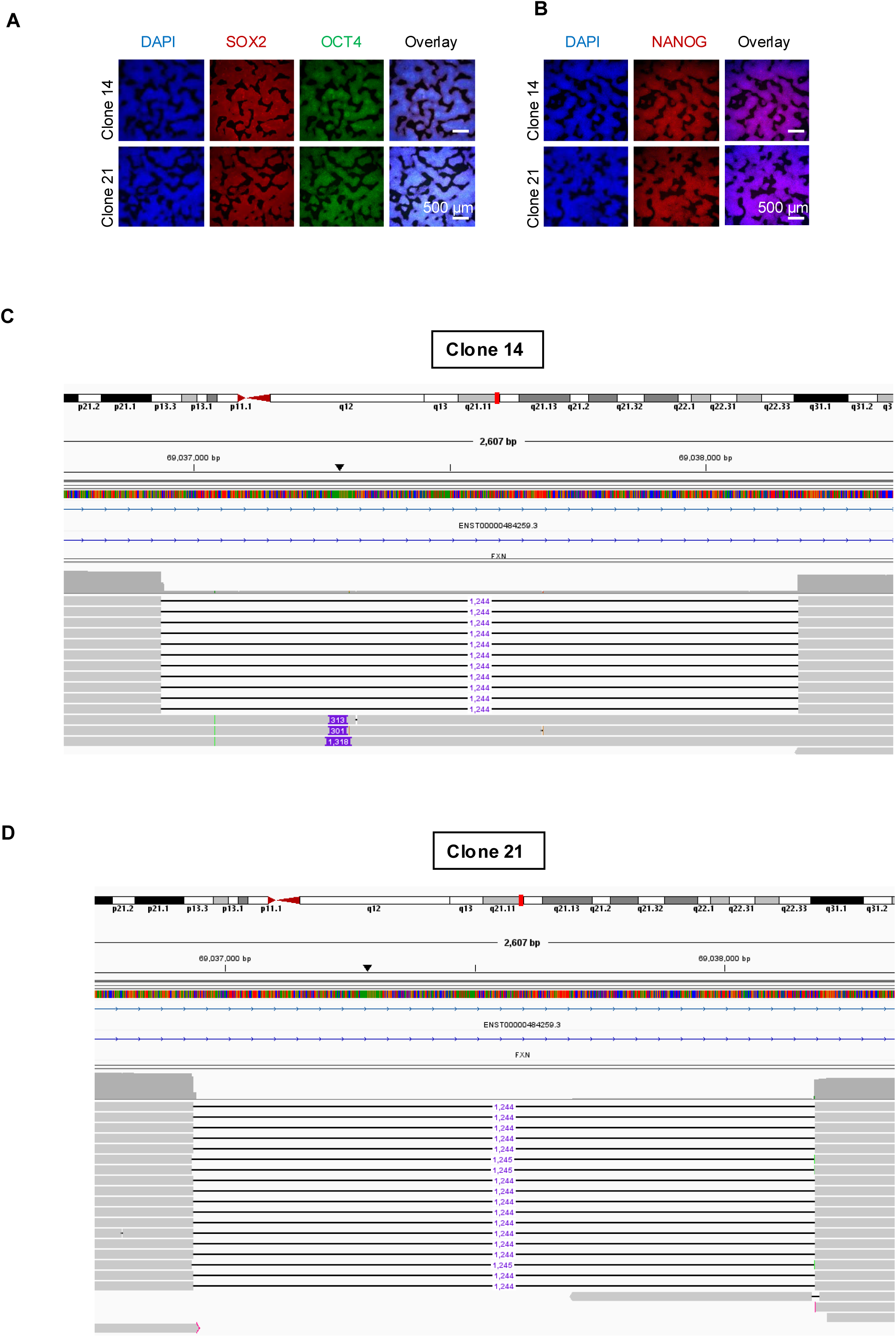
Characterization of isogenic gene-corrected FRDA iPSC lines. (A) Expression of pluripotency markers in two established cell clones of gene-corrected FRDA iPSC lines (GM23913). (B-C) Visualization of FXN intron 1 targeted locus with integrated genome viewer (IGV) in the gene-corrected isogenic iPSCs from a FRDA patient (GM23913). PacBio long-read sequencing of the iPSC clones confirms excision in the FXN intron 1 locus (1244 bp + GAA repeats).

**Figure S6.**
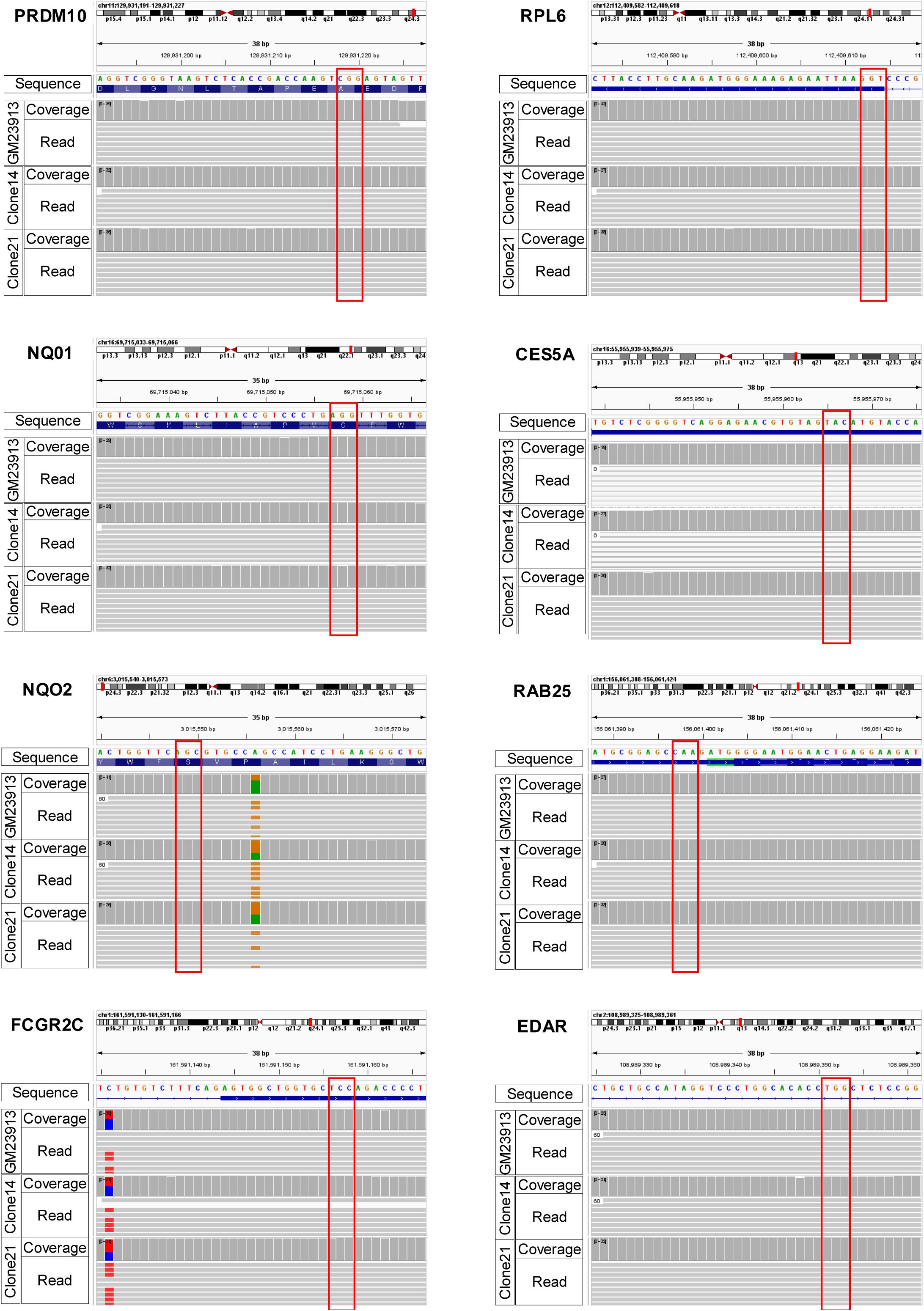
Analysis of the 4 highest ranked off-target genomic sites in the corrected, isogenic GM23913 iPSC clones. For 5’ and 3’ gRNA utilized for the genomic deletion, the corrected isogenic GM23913 iPSC clones showed no off-target deletions in any of the predicted off-target sites. The PAM sequences are highlighted in the red box.

**Figure S7.**
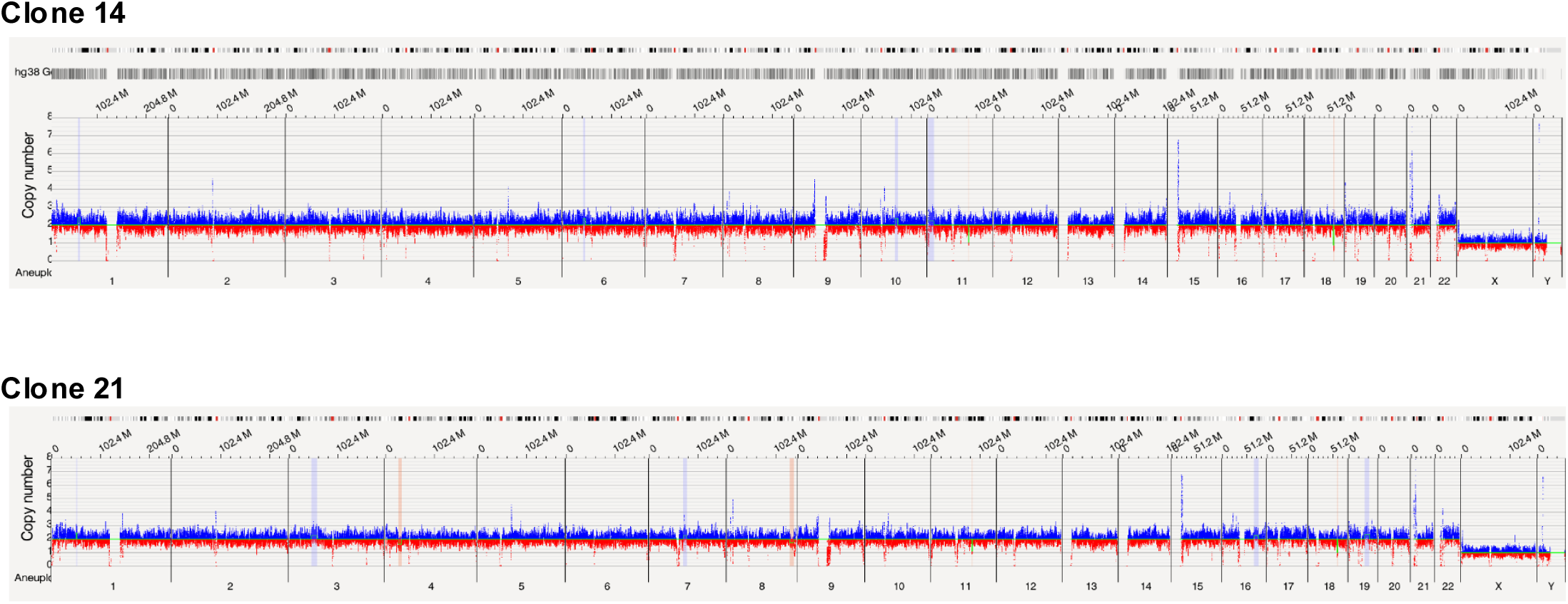
Copy number frequency plots of the corrected, isogenic GM23913 iPSC clones identified by optical genome mapping. Plots showing typical copy number frequency through the entire chromosomes, i.e. 2 copy numbers in chromosomes 1-22 and 1 copy numbers in chromosome x and y, respectively.

