## Supplementary materials for "Modeling Friedreich’s ataxia with Bergmann glia-enriched human cerebellar organoids"

**Supplementary Table 1.** List of primary and secondary antibodies.

| Type | Antibody | Vendor | Catalog Number | Application | Dilution Factor |
|---|---|---|---|---|---|
| Primary | WNT1 | Santa Cruz | sc-514531 | Western blot | 1:20 |
| Primary | FGF8 | Santa Cruz | sc-293479 | Western blot | 1:20 |
| Primary | EN1 | Santa Cruz | sc-398534 | Western blot | 1:20 |
| Primary | EN2 | Santa Cruz | sc29331 | Western blot | 1:20 |
| Primary | Hoxa2 | Novus Biologicals | nbp2-58865 | Western blot | 1:20 |
| Primary | GAPDH | Santa Cruz | sc-3765062 | Western blot | 1:10000 |
| Primary | SOX2 | Abcam | ab79351 | Immunostaining | 1:100 |
| Primary | SOX2 | Millipore | ab5603 | Immunostaining | 1:100 |
| Primary | TUJ1 | Biolegend | 801202 | Immunostaining | 1:200 |
| Primary | Kirrel2 | R&D Systems | mab2564 | Immunostaining, western blot | 1:100, 1:20 |
| Primary | SKOR2 | Atlas Antibodies | HPA046206 | Immunostaining, western blot | 1:100, 1:20 |
| Primary | PAX2 | Biolegend | 901001 | Immunostaining, western blot | 1:100, 1:20 |
| Primary | Barhl1 | Novus Biologicals | NBP1-86513 | Immunostaining, western blot | 1:100, 1:20 |
| Primary | S100β | Abcam | ab52642 | Immunostaining, western blot | 1:100,1:20 |
| Primary | ATOH1 | ABclonal | A11477 | Immunostaining | 1:100 |
| Primary | ALDH1L1 | Cell Signaling Technologies | CST85828 | western blot | 1:20 |
| Primary | EAAT1 | Cell Signaling Technologies | CST5684 | western blot | 1:20 |
| Primary | SOX9 | Abcam | ab185230 | western blot | 1:20 |
| Primary | Calbindin | Abcam | ab75524 | Immunostaining, western blot | 1:500, 1:20 |
| Primary | GABA | Sigma | A2052 | Immunostaining | 1:100 |
| Primary | NeuN | Abcam | ab104224 | Immunostaining | 1:100 |
| Primary | NeuN | Abcam | ab177487 | Immunostaining, western blot | 1:200, 1:50 |
| Primary | SMI32 | Biolegend | 801701 | Immunostaining | 1:100 |
| Primary | TBR1 | Abcam | ab183032 | Immunostaining, western blot | 1:100, 1:20 |
| Primary | Neurogranin | Millipore | ab5620 | Immunostaining, western blot | 1:200, 1:50 |
| Primary | MAP2 | Millipore | mab3418 | Immunostaining | 1:200 |
| Primary | GFAP | Dako | z0334 | Immunostaining, western blot | 1:500, 1:20 |
| Primary | OCT4 | Santa Cruz | sc-5279 | Immunostaining | 1:200 |
| Primary | Nanog | Cell Signaling Technologies | 4903s | Immunostaining | 1:200 |
| Primary | SOX2 | R&D Systems | AF2018 | Immunostaining | 1:100 |
| Primary | GBX2 | Abnova | H00002637 | Immunostaining | 1:50 |
| Primary | PAX6 | Biolegend | 862002 | Immunostaining | 1:100 |
| Primary | GBX2 | R&D Systems | af4638 | Western blot | 1:20 |
| Primary | Frataxin | Abcam | ab110328 | Western blot | 1:20 |
| Primary | Cleaved caspase-3 | Cell Signaling Technologies | 9664s | Western blot | 1:50 |
| Primary | Pro-caspase-3 | Cell Signaling Technologies | 9665s | Western blot | 1:50 |
| Primary | aconitase 2 | Abcam | ab110321 | Western blot | 1:20 |
| Primary | Acetyl H4K12 | Cell Signaling Technologies | CST13944 | Western blot | 1:20 |
| Primary | Acetyl H4K5 | Cell Signaling Technologies | CST8647 | Western blot | 1:20 |
| Primary | Histone4 | Cell Signaling Technologies | CST2935 | Western blot | 1:20 |
| Primary | DRP1 | Cell Signaling Technologies | 8570 | Western blot | 1:20 |
| Primary | P-DRP1 (Ser637) | Cell Signaling Technologies | 6319 | Western blot | 1:20 |
| Primary | MFF | Cell Signaling Technologies | 84580 | Western blot | 1:20 |
| Primary | Mitofusin-1 | Cell Signaling Technologies | 14739 | Western blot | 1:20 |
| Primary | Mtofusin-2 | Cell Signaling Technologies | 11925 | Western blot | 1:20 |
| Primary | OPA1 | Cell Signaling Technologies | 80471 | Western blot | 1:20 |
| Primary | TOM20 | Cell Signaling Technologies | 42406 | Western blot | 1:20 |
| Secondary | Secondary | 1:500 | Thermo Fisher Scientific | Donkey anti-mouse Alexa 488 | A-21202 |
| Secondary | Secondary | 1:500 | Thermo Fisher Scientific | Donkey anti-rabbit Alexa 568 | A-10042 |
| Secondary | Secondary | 1:500 | Thermo Fisher Scientific | Donkey anti-goat Alexa 647 | A-21447 |
| Secondary | Secondary | 1:500 | Thermo Fisher Scientific | Donkey anti-mouse Alexa 568 | A-10037 |
| Secondary | Secondary | 1:500 | Thermo Fisher Scientific | Donkey anti-rabbit Alexa 488 | A-21206 |

**Supplementary Table 2.** List of primer sequences.

| **Target** | **Forward** | **Reverse** |
|---|---|---|
| **FXN intron1 GAA expansion** | GAGGTCTAACCTCTAGCTGCTC | AAGCCCAATACGTGGCAG |
| **FXN CRISPR-Cas9 validation** | GGAGGGAACCGTCTGGGCAAAGG | CAATCCAG GACAGTCAGGGCTTT |

**Supplementary Table 3.** Guide RNA sequences.

| gRNA | Sequences |
|---|---|
| **gRNA1 (up)** | GG CGU ACC AGC CAC UCU GAA GUU UUA GAG CUA UGC U |
| **gRNA2 (down)** | CA AGA UGU GCA AGG GAA CUA GUU UUA GAG CUA UGC U |

### Methods

#### Cell culture and organoid formation

Human induced pluripotent stem cells (iPSCs) were obtained from NIGMS Human Genetic Cell Repository at the Coriell Institute for Medical Research (GM25256, GM23404, GM23903) and NIH Common Fund (NCRM5). Human embryonic stem cells (hESCs) were obtained from WiCell Research Institute (WA09). Human pluripotent stem cells (hPSCs) were maintained in feeder-free conditions using mTeSR™ Plus (STEMCELL Technologies) and vitronectin (VN)-coated plates (Thermo Fisher Scientific). Cell lines were confirmed to be karyotypically normal and mycoplasma-free (MycoAlert Detection Kit, Lonza). Cells were routinely passaged when cultures reached 70-90% confluency (every 3 to 4 days) using 0.5 mM EDTA diluted in phosphate buffered saline (PBS) without calcium or magnesium (Thermo Fisher Scientific). Cell cultures were maintained at 37°C under humidified 5% $CO_2$ and atmospheric $O_2$.

To generate human cerebellar organoids (hCBOs), hPSCs were detached using Accutase (Thermo Fisher Scientific) and plated in 24-well AggreWell™ 800 plates (STEMCELL Technologies) at a density of 3,000-6000 cells per cell aggregate, i.e. EB, in 2mL/well of growth factor-free chemically defined media (gfCDM), which is modified from a previous study consisting of DMEM/F12 GlutaMAX™ (Gibco), chemically defined lipid concentrate (1% v/v, Life Technologies), monothioglycerol (450 µM, Sigma), apo-transferrin (15 µg/ml, Sigma), bovine serum albumin (5 mg/ml, Sigma), and supplemented with insulin (7 µg/ml, Sigma).[1] CEPT cocktail (50 nM chroman 1 (MedChem Express), 5 µM emricasan (Selleckchem), 1X polyamine supplement (Sigma), and 0.7 µM trans-ISRIB (Tocris)) was used in the first 2 days of cell aggregation. FGF2 (50 ng/ml, R&D Systems), A83-01(2 µM, Tocris), and LDN-193189 (100 nM, Tocris) were added on day 0.[2, 3] CHIR99021 (0.5 µM, Tocris) and recombinant human FGF2 (50 ng/ml, R&D Systems) were added on day 2. The entire medium was replaced on day 2 and every other day with gfCDM supplemented with A83-01, LDN-193189, CHIR99021, and FGF2. On day 7, floating aggregates were transferred to ultra-low attachment 6-well plates (Corning) at one well of a 24-well plate to one well of a 6-well plate. Aggregates were cultured in 4 mL/well of gfCDM media with 2/3 of the initial amount of A83-01, LDN-193189, CHIR99021, and FGF2 supplements on day 7 and 1/3 on day 10. The plates were placed on an orbital shaker starting on day 10. From day 14 to day 21, media was replaced with gfCDM supplemented with recombinant human FGF19 (100 ng/ml, R&D systems). From day 14 onward, the entire medium was replaced every 3-4 days. From day 21 to day 35, the formed organoids were cultured in DMEM/F12 GlutaMAX™ (Gibco), chemically defined lipid concentrate (1% v/v, Life Technologies), N2 supplement (Gibco), and B27 minus vitamin A (Gibco) as the base media with the addition of glial induction factors (consisting of Delta-like protein-1 (DLL1), Jagged-1, LIF, Oncostatin M, and ciliary neurotrophic factor 1 (CNTF), 10 ng/mL each, R&D Systems).[4] Recombinant human SDF1 (300 ng/ml, R&D systems) was added to the culture from day 28 to day 35. From day 35 onward, the media was replaced with final maturation media that consisted of DMEM/F12 GlutaMAX™ (Gibco), chemically defined lipid concentrate (1% v/v, Life Technologies), N2 supplement (Gibco), B27 (Gibco), recombinant human BDNF (50 ng/mL, R&D systems), GDNF (50 ng/mL, R&D systems), NT3 (50 ng/mL, R&D systems), CNTF (10 ng/mL, R&D systems), and T3 (1 µM, Tocris). The medium was replaced every 3-4 days. For the CHIR99021 dose response studies, hCBOs were cultured using the same protocol up to day 14, but with various CHIR99021 concentrations at either 0, 0.5, 1, or 2 µM starting at day 2. At day 14, the formed organoids were analyzed by Western blot.

To capture dendritic arborization of Purkinje cells, organoids were dissociated at day 60 using an Embryoid Body Dissociation Kit (Miltenyi Biotech) and gentleMACS Dissociator (Miltenyi Biotech) in a gentleMACS C Tube (Miltenyi Biotech) following the manufacturer's protocol. The cell suspension was filtered through a 70-µm strainer (Miltenyi Biotech) to remove cell debris and clumps. The strained cell suspension was centrifuged at 300 g for 5 min and plated on wells coated with poly-L-ornithine (15 µg/mL, Sigma) and Laminin (20 µg/mL, Sigma), at a seeding density of 100,000 cells/cm$^2$. Cells were cultured with the maturation media which was replaced every 3-4 days. 30 days after re-plated, cells were fixed in 4% paraformaldehyde (PFA; Thermo Fisher Scientific) for 20 min at RT and washed three times with PBS.

To generate cortical organoids, we implemented a modified method from a previously published protocol.[5] iPSCs were dissociated into single cells using Accutase (Thermo Fisher Scientific) and plated in 96-well ULA round-bottom plates (Corning) at a density of 6,000 cells per well with the CEPT cocktail in cortical differentiation medium (CDM) I, containing Glasglow-MEM (Gibco), 20% Knockout Serum Replacement (Gibco), 0.1 mM Minimum Essential Medium non-essential amino acids (Gibco), 1 mM Sodium Pyruvate (Gibco), and 0.1 mM 2-mercaptoethanol (Gibco). TGFβ inhibitor SB431542 (Tocris) and WNT inhibitor IWR1 (Tocris) were added at a concentration of 5 μM and 3 μM, respectively. Media was changed every 3 days with CDM1 48 h after seeding. After 18 days, the aggregates were cultured in 100-mm ULA culture dishes (Corning) on an orbital shaker and media was changed to CDM II containing DMEM/F12 medium (Gibco), 2 mM Glutamax (Gibco), 1% N2 (Gibco), and 1% Chemically Defined Lipid Concentrate (Gibco). Media was changed every 3 days until day 35.

#### Western blot analysis

Organoids were washed in PBS, resuspended in RIPA buffer (Thermo Fisher Scientific) supplemented with half protease inhibitor cocktail (Thermo Fisher Scientific) and lysed by sonification. Cell debris was removed by centrifugation at 14,000g for 15 min. Protein quantification was performed using the BCA protein assay kit (Thermo Fisher Scientific). The Wes and Jes automated western blotting systems (ProteinSimple) were utilized following the manufacturer's instructions. All western blot data are presented by lanes in virtual blot-like images. For western blot quantification, band intensity was calculated using ImageJ. The relative amount of the target protein was normalized to GAPDH. Detailed information on primary and secondary antibodies is provided in Supplementary Table 1.

#### Histological analysis

Organoids were fixed in 4% PFA at RT for 3 h, followed by three washes with PBS. Subsequently, they were incubated in 30% sucrose at 4°C overnight. Tissues were then embedded in 7.5% gelatin/10% sucrose embedding solution (Sigma), molded in dry ice/acetone solution, sectioned into 10-μm slices, and mounted onto microscope slides (Fisher Scientific) for staining. For histological analysis, sections underwent staining with H&E, dehydration in a series of ethanol concentrations (70%, 80%, 95%, 100%) and 100% xylene, stained in hematoxylin and eosin dye, and mounted on coverslips using Permount mounting medium (Fisher Scientific). For immunohistochemical analysis, sections were permeabilized and blocked with 0.3% Triton X-100 and 5% BSA in PBS for 1 h. Detailed information regarding primary and secondary antibodies can be found in Supplementary Table 1. Slides were mounted using ProLong Glass Antifade Mountant with NucBlue Stain (Thermo Fisher Scientific), and fluorescence images were captured either using the Akoya PhenoImager HT or Zeiss LSM 710 confocal microscope with appropriate filters.

#### Image-based neuronal migration analysis

$NeuN^+/GFAP^+/DAPI^+$ and $Calbindin^+/NeuN^+/DAPI$ images captured using the Zeiss LSM 710 confocal microscope were analyzed using Simple Linear Iterative Clustering algorithm, a superpixel algorithm that clusters pixels based on color similarity and proximity in the image plane, and Graph Cut to quantify neuronal migration in cerebellum organoids. This approach enabled the precise measurement of neuronal migration dynamics within the organoid structure. A deep learning method in Cellpose was employed to individually segment NeuN, GFAP, Calbindin, and DAPI within the hBCO. Utilizing the superpixel machine learning algorithm, precise measurements were made for the positions of calbindin/NeuN and the boundaries of the GFAP/NeuN layer. At the final stage, the obtained results in pixels were converted into μM units.

**Microelectrode array (MEA) analysis for electrophysiology study**

Neuronal activity was analyzed using the Maestro Pro multiwell microelectrode array system (Axion Biosystems) according to the manufacturer's protocol. Briefly, day 60 hCBOs were plated on PLO/Laminin coated 48-well MEA plates following the manufacturer's protocol with final maturation media. The neuronal activity was recorded for 5 min on day 80, which is 2-3 weeks post plating. For ATP stimulation, the neuronal activity was recorded immediately following the addition of 100 μM ATP (Sigma).

**Sanger sequencing, PCR gel electrophysiology, and PacBio long sequencing**

Human iPSCs were washed in PBS 1X, centrifuged, and processed for genomic DNA extraction. Genomic DNA was performed with GeneJet genomic DNA purification column kit (Thermo Fisher Scientific) according to the manufacturer's protocol. For PCR amplification, Platinum™ SuperFI II PCR Master Mix (Thermo Fisher Scientific) was used following the manufacturer's protocol. Primer sets can be found in Supplementary Table 2. The amplified PCR samples were run on E-Gel™ 1% agarose gels with SYBR™ Safe DNA Gel Stain (Invitrogen) for visualization with E-Gel™ sample loading buffer (Invitrogen). For Sanger sequencing, primer extension sequencing was performed by AZENTA Life Sciences using commercial dye terminators. The reactions were then run on an ABI 3730xl DNA Analyzer (Applied Biosystems). For PacBio long sequencing, the short fragments (<10kb) of the HMW gDNA from the 5 hiPSC samples were removed using Short Read Eliminator (SRE) kit (Pacific Biosciences). 2 μg of DNA were sheared to 14-19 kb using the Megaruptor 3 (Diagenode, Inc.) at shear speed 31. The sheared DNA were purified using 1X SMRTbell cleanup beads and used to prepare whole genome libraries using SMRTbell Prep Kit 3.0 (Pacific Biosciences). Sequencing primer was annealed and Revio polymerase was bound to the library prior to loading using the Revio Polymerase kit (Pacific Biosciences). Each library was loaded on one Revio SMRTcell at 200 pM loading concentration. Sequencing was performed with a 24h movie time on Revio.

**ROS quantification**

For ROS quantification, day 60 hCBOs were washed twice in DPBS (Thermo Fisher Scientific) and dissociated into a single-cell suspension using Embryoid Body Dissociation Kit (Miltenyi Biotech) and gentleMACS Dissociator (Miltenyi Biotech) in a gentleMACS C Tube (Miltenyi Biotech) following the manufacturer's protocol. The cell suspension was filtered through a 70-μm strainer (Miltenyi Biotech) to remove cell debris and clumps. The strained cell suspension was centrifuged at 300 g for 5 min and stained with $H_2$DCFDA reagent (Thermo Fisher Scientific) according to the manufacturer's protocol. The stained cells were analyzed by flow cytometry (SONY SH800) using the FITC filter.

**Electron microscopy**

Day 60 hCBOs were first fixed in 2% glutaraldehyde (v/v) in sodium cacodylate buffer (0.1 M, pH 7.4), followed by rinsing in cacodylate buffer, and post-fixation in 1% osmium tetroxide (v/v) solution (Electron Microscopy Sciences) in cacodylate buffer. After further rinsing in cacodylate buffer and acetate buffer (0.1 M, pH 4.0), the organoids underwent uranyl acetate (0.5% w/v, pH 4.5) en bloc staining. Dehydration was carried out using a series of ethanol concentrations (e.g., 35%, 50%, 75%, 95%, and 100%), followed by washing and overnight exposure to pure epoxy resin (Electron Microscopy Sciences). The next day, the organoids were embedded in fresh epoxy resin and processed further in a 55°C oven for 48 h. The cured epoxy was separated by submerging it in liquid nitrogen. Thin sections (60-70 nm) were then prepared using a diamond knife and ultramicrotome (Leica). These sections were mounted onto 150 copper mesh grids and counter-stained with aqueous uranyl acetate (0.5% w/v) and Reynold's lead citrate. Finally, the samples were examined using an electron microscope (Hitachi), with images captured using a CCD camera (AMT).

#### RNA bulk-sequencing analysis

RNA was extracted (three samples for each group, each sample consisting of three organoids) using the RNeasy Mini Kit (Qiagen). RNA was quantified using the Agilent RNA ScreenTape System (Agilent) on a 4200 TapeStation (Agilent). RNA (500ng, RIN >8) was used to prepare RNA-seq libraries with the KAPA mRNA HyperPrep Kit (Roche) according to the manufacturer's protocol. Sequencing libraries were quantified by qPCR using the KAPA library quantification kit (Roche) using QuantStudio 12K Flex Real-Time PCR System (Thermo Fisher Scientific). All samples were normalized according to concentration and pooled. Libraries were loaded and sequenced using the Illumina Novaseq 6000 system. Bioinformatics analysis for bulk RNA-seq was carried out using the computational resources of the NIH HPC Biowulf cluster (http://hpc.nih.gov) using R language 3.6.0 (https://cran.r-project.org/). Bulk RNA-Seq samples were quality trimmed using Trimmomatic 0.36 and the TruSeq3 paired-end adapters. STAR aligner 2.7.6a followed by HTSeq-count 0.9.1 produced deduplicated counts of reads in genes. UCSC gene counts were normalized using the default median-of-ratios method in DESeq2 1.24.0. Differential expression tests used the lfcShrink function and gene set enrichment was performed using Enrichr API package enrichR 1.0.

#### HDAC inhibitor testing on day 60 hCBOs

GM23913 hCBOs at day 60 were either untreated or treated with HDAC inhibitors at their reported $IC_{50}$ value for HDAC inhibition, i.e. BML-210, SBHA, SAHA all at 5 µM, TSA at 0.1 µM, and Valproic acid at 400 µM.[6] The organoids were analyzed by western blot after 24 h of treatment.

#### Generation of corrected, isogenic FRDA hiPSC lines

CRISPR RNA pairs were designed in regions of intron 1 of the FXN gene (Gene ID:2395) with CRISPOR http://crispor.tefor.net, which has been previously reported (see Supplementary Table 3 for sequences).[7] Next, gRNA was formed by thermal incubation of 100 µM crRNA, 100 µM ATTO™ 550-tagged tracrRNA, and nuclease-free duplex buffer (all from Integrated DNA Technologies) at a ratio of 4.4:4.4:1.2 at 95°C for 5 min followed by incubation at RT for 10 min. Next, the ribonucleoprotein complex was formed by incubating the formed gRNA with 36 µM Alt-R Cas9 nuclease (Integrated DNA Technologies) at a ratio of 1:1 in RT for 20 min. To generate corrected, isogenic hiPSC lines, hiPSCs were electroporated using the NEON Transfection system (Thermo Fisher Scientific) according to the manufacturer's protocol. In brief, cells were dissociated into single cells with Accutase (Thermo Fisher Scientific), resuspended in Neon Resuspension Buffer to a density of $2.2x10^7$ cells/mL, and mixed with the formed RNP complexes at a ratio of 1 (RNP complex of gRNA1):1 (RNP complex of gRNA2):9 (cell suspension). The mixture was electroporated using 10 µl-pipette tips (1400V, 20ms, 1 pulse). Electroporated cells were seeded into VN-coated 24-well plates containing mTeSR™ Plus supplemented with CEPT. 3 days after electroporation, cells were detached with Accutase and either analyzed with PCR gel electrophoresis to confirm the designed CRISPR-Cas9 system efficacy or printed with Hana (Namocell) single-cell printer for clonal expansion following a previously published protocol.[8]

#### Optical genomic mapping

Ultra-high molecular weight (UHMW) gDNA was extracted with the Bionano Prep SP-G2 Blood & Cell DNA Isolation Kit (Bionano Genomics) as described in the Bionano Prep SP Frozen Cell Pellet DNA Isolation Protocol (Bionano Genomics). Briefly, 1.5 million frozen pelleted cells were thawed in a 37°C bath and resuspended in DNA Stabilizing Buffer (Bionano Genomics). After that, the cells were lysed in the presence of detergents, proteinase K, and RNase A. The UHMW gDNA was bound to a silica Nanobind Disk (Bionano Genomics), washed, and eluted. The extracted gDNA was equilibrated for 10 days at 4°C to homogenize and then quantified with Qubit dsDNA BR assay kit (Thermo Fisher Scientific). The isolated UHMW gDNA was fluorescently labeled using the Bionano Prep DLS-G2 labeling kit (Bionano Genomics, 80046) according to the Bionano Prep DLS-G2 Protocol (Bionano Genomics). In short, Direct Label Enzyme (DLE-1) and DL-green fluorophores were used to label 750 ng of purified gDNA at a specific sequence motif. After a cleanup of the fluorophores excess, DNA backbone was counterstained overnight before quantitation with Qubit dsDNA HS assay kit (Thermo Fisher Scientific). Finally, the labeled UHMW gDNA molecules were loaded on a Saphyr G3.3 chip for sequential imaging across nanochannels on the Saphyr instrument (Bionano Genomics). Genome analysis was performed using Bionano Solve-v 3.7 and Access 1.7.2. For genome assembly and variant analysis, Bionano De novo assembly pipeline was run to assemble a genome from the molecule files (*bnx) for each sample with human GRCh38 reference provided.

**Statistical analysis**

All results are shown as mean ± SEM. Statistical tests included unpaired, two-tailed Student's t-tests and one-way ANOVA for multiple comparisons with Tukey's significant difference post hoc test using GraphPad Prism 9.0.0. A $P$ value of $<0.05$ was considered statistically significant. $P$ values denote as $^*p<0.05$, $^{**}p<0.01$, $^{***}p<0.001$, or $^{****}p<0.0001$.
